# Protein-free membrane fusion: a refined view of the delicate fusogenic properties of calcium

**DOI:** 10.1101/2025.11.19.689405

**Authors:** M. Accorsi, R. Ní Earchaí, N. Yandrapalli, S. Pramanik, R. Dimova

## Abstract

In the late 20th century, calcium took on the identity of an independent fusogen, when it was found to induce fusion of anionic large unilamellar vesicles (LUVs), yet its ability to drive fusion in cell-sized membranes remains poorly understood. Here, we directly quantify calcium-mediated fusion of giant unilamellar vesicles (GUVs) using a microfluidic trapping platform combined with confocal microscopy, enabling simultaneous measurement of lipid mixing, content mixing, and fusion outcomes across hundreds of single vesicles. We systematically map fusion efficiency as a function of calcium concentration, membrane composition, and mechanically imposed tension. We find that calcium-induced fusion of GUVs in the absence of proteins is remarkably fickle and composition-sensitive, as the vesicles need to be sufficiently instable to allow the opening of the fusion pore, yet stable enough to prevent bursting and collapse. Negatively charged GUVs containing high fractions of DOPE exhibit the highest fusogenic responsiveness, whereas other compositions undergo extensive lipid mixing without pore formation. Increasing membrane tension can shift this balance and promote full fusion, revealing a narrow parameter space in which calcium acts as an effective protein-free fusogen for cell-sized membranes. These findings clarify longstanding discrepancies between LUV- and GUV-based calcium fusion assays and provide quantitative design rules for employing calcium as a fusogen in synthetic biology and membrane-reconstitution studies, where controlled membrane growth, vesicle-vesicle fusion, and module integration are central to building and sustaining artificial cells.

**SIGNIFICANCE:** Calcium is widely regarded as a potent, protein-free fusogen in nanometer-scale lipid vesicles, yet its relevance for fusion between cell-sized membranes remains unresolved. This gap limits our ability to translate decades of LUV-based fusion studies to more physiologically realistic systems and restricts the use of calcium in synthetic cell engineering. By directly quantifying calcium-mediated fusion in giant unilamellar vesicles (GUVs), we identify the membrane compositions and mechanical conditions under which calcium can reliably drive full fusion rather than mere lipid mixing. Our findings provide the first systematic map of calcium’s fusogenic parameter space in cell-sized membranes, enabling more informed design of fusion-based assays, reconstitution experiments, and strategies for membrane growth in artificial cells.

## 1 INTRODUCTION

Cell fusion has long been the object of extensive research due to its crucial role in various biological processes, including viral infection mechanisms (1-8), cell fertilization (9-11) and cell signaling (12-15). Elucidating the steps and molecular players behind fusion is essential for advancing our knowledge across multiple areas of biology and medicine (1,13,16,17). Understanding cell fusion may inform us about disease mechanisms, while knowledge of viral fusion can provide tools to counter infections, blocking the fusion process such as in the case of HIV (1,5-8).

Membrane fusion is typically triggered and carried out by fusogens, which include both proteins, such as viral fusion proteins and peptides, and non-protein molecules such as specific lipids and calcium ions. While protein fusogens drive fusion through conformational changes (18,19), non-protein fusogens such as calcium promote membrane merging by interacting with lipid headgroups (20-22). In addition to this direct role, calcium can also act as a trigger for the activating fusion proteins in biological processes (4,14,19,23). Thus, calcium emerges as a key player in both protein-dependent and protein-free fusion models (20,21,24-27).

Much of the research carried out in the biochemical and medical fields has focused on fusion proteins. Indeed, a complex array of protein machinery is often involved in triggering fusion, requiring the delicate coordination of multiple interacting partners, such as proteins and cell signals, to correctly carry out the fusion process. However, the lipid membrane (the very platform on which these proteins act) has received comparatively less attention. Evidence suggests that lipid microdomains at the fusion site play an active regulatory role, either by locally reorganizing to facilitate pore formation or by modulating palmitoylated proteins (21,28-33). A deeper understanding of membrane composition and mechanics is also proving invaluable in applied fields such as vaccine development (34) and targeted drug delivery (35), both of which are heavily dependent on controlled membrane fusion. Furthermore, lipid vesicles as membrane models have long served as a means for micro-compartmentalization and transport (36).

Because membrane curvature plays a fundamental role in fusion (30,37-39), the local curvature of lipid vesicles likely influences specific steps in the fusion process (39-41). Interestingly, the asymmetric distribution of calcium across membrane leaflets is proposed to induce curvature, though experimental findings remain debated. Some studies reports that binding of calcium to negatively charged membranes induces positive spontaneous curvature (42), while others observe the opposite effect – generation of negative spontaneous curvature (43,44). The latter observation aligns with simulations (20), lending further credibility to calcium’s role as an active fusogen. Importantly, large absolute values of spontaneous curvature correspond to substantial spontaneous (or intrinsic) membrane tension (45), which resists further deformation. Thus, changes in spontaneous curvature not only reshape the local membrane landscape but also alter the mechanical stress balance across the bilayer, modulating both the likelihood and the pathway of fusion (41,46).

Various model membrane systems have been developed to study fusion, each with advantages and limitations (40). These include reconstituted protein systems, which allow precise control over protein composition but may suffer from complexity and low efficiency. Supported lipid bilayers enable high-resolution imaging but may be affected by substrate interactions. Large unilamellar vesicles (LUVs) are easy to prepare and suitable for bulk measurements, but their small size represents a highly curved environment and limits direct observation. Indeed, in the second half of the 20th century, many studies employed LUVs to investigate the fusogenic properties of calcium, highlighting its ability to induce fusion between negatively charged vesicles in the absence of proteins (26-28,47,48). However, the interpretation of such bulk assays can be problematic, as transient rupture and resealing events, which are undetectable in ensemble measurements, may mimic fusion signals, leading to potential overestimation of true fusion efficiency.

In the context of synthetic biology, membrane fusion is increasingly viewed as a powerful mechanism for driving controlled growth, communication, and material exchange in artificial cells (49). Fusion-mediated lipid and protein delivery offers a route for programmed expansion of vesicle membranes (50), while regulated content mixing enables stepwise assembly of biochemical pathways within compartmentalized reaction networks (51-53). Minimal cells that grow, divide, or communicate through engineered fusion events represent an emerging frontier in bottom-up biology (54), where understanding how membrane composition and physical parameters modulate fusion efficiency becomes essential. Calcium-mediated fusion between lipid vesicles therefore provides not only a model for fundamental biophysical processes but also a potential tool for constructing life-like systems capable of adaptive remodeling and growth.

In recent years, giant unilamellar vesicles (GUVs) have gained popularity as membrane models (55,56). Their micrometer-scale size closely resembles that of cells, making them a more physiologically relevant alternative to LUVs. One of their key advantages is the ability to directly observe individual vesicles over the course of an experiment using optical microscopy. This facilitates the study of individual stages of the fusion process and eliminates the need for indirect bulk measurements or super-resolution techniques. Moreover, GUVs can be directly subjected to controlled mechanical perturbations, such as micromanipulation, electrodeformation, or osmotic gradients, allowing the quantitative assessment of membrane tension and its influence on fusion dynamics. GUVs have been widely used to study fusion mediated by various fusogens, including SNARE proteins (57,58), electric fields (59,60), ligands (12,61), nanoparticles (25), and oppositely charged lipids (36,62-64) to mention a few. Their versatility makes them an ideal platform for investigating calcium-mediated membrane fusion.

Much of the existing research on calcium-induced membrane fusion has been conducted using LUVs (26-28,36,47). This approach, however, leaves several important questions unanswered. Could findings on LUV systems be directly translated to membranes with quasi-zero curvature, such as those in GUV-GUV and cell-cell fusion? How does calcium-mediated fusion efficiency vary with membrane compositions? Are the fusion mechanisms observed in highly curved small and large unilamellar vesicles (SUVs and LUVs) applicable to larger, less curved membranes such as GUVs or biological cells? Moreover, how reliable are the results of LUV-based fusion assays when applied to GUVs? Addressing these questions could provide valuable insights for assessing the broader applicability of calcium as a fusogen and its role in biological membrane fusion processes.

With this in mind, here we employ microfluidics and microscopy-based approaches to directly observe and test the effect of calcium on membrane fusion; first by analyzing the effect of increasing calcium concentrations comparable to those used in previous in vitro studies (26-28,36,47) rather than in the lower concentrations found in biological systems (65), then by exploring GUVs of varying lipid compositions, and finally by observing the role that membrane tension plays. We also propose a versatile framework for quantifying hemifusion and full fusion across large populations of GUVs, combining multiple microscopy techniques, including Förster resonance energy transfer (FRET) and direct optical observation, to capture both lipid and content mixing events. This approach allows systematic, high-resolution analysis of fusion dynamics at the single-vesicle level.

## 2 MATERIALS AND METHODS

### 2.1 Materials

The phospholipids 1-palmitoyl-2-oleoyl-sn-glycero-3-phosphocholine (POPC), 1-palmitoyl-2-oleoyl-sn-glycero-3-phospho-L-serine (sodium salt) (POPS), 1,2-dioleoyl-sn-glycero-3-phosphoethanolamine (DOPE), 1,2-dioleoyl-sn-glycero-3-phosphocholine (DOPC) and the fluorescent dyes 1,2-dipalmitoyl-sn-glycero-3-phosphoethanolamine-N-(lissamine rhodamine B sulfonyl) (ammonium salt) (Rh-DPPE) and 1,2-dioleoyl-sn-glycero-3-phosphoethanolamine-N-(7-nitro-2-1,3-benzoxadiazol-4-yl) (ammonium salt) (NBD-DOPE) were purchased from Sigma-Aldrich Chemie GmbH, Germany. The lipids 1,2-dioleoyl-sn-glycero-3-phosphoethanolamine labeled with Atto 633 (Atto633-DOPE) and with Atto 488 (Atto488-DOPE) were purchased from ATTO-TEC GmbH North Rhine-Westphalia, Germany.

Lipid solutions were prepared in chloroform and stored at -20 °C until use. Sucrose, glucose, bovine serum albumin (BSA) and calcium chloride (CaCl_2_) were purchased from Sigma-Aldrich and used as received. SYLGARD 184 elastomer kit was purchased from Biesterfeld Spezialchemie GmbH, Hamburg, Germany.

### 2.2 Vesicle preparation

#### 2.2.1 GUV preparation

We employed an electroformation protocol for the preparation of GUVs. Due to the difficulty in growing GUVs with high amounts of DOPE, the usual electroformation settings where modified according to the parameters specified in (48). For each GUV formation, two indium tin oxide (ITO)-coated glass plates were used (Praezisions Glas & Optik, Iserlohn, Germany). Small amounts (10 μL) of 3 mM lipid in chloroform solution were spread on each of the plates. The lipid-covered glasses were then left in a vacuum desiccator for a minimum of one hour to evaporate the chloroform. After drying, the two plates and a rectangular Teflon frame were assembled to form an electroformation chamber. 2 mL of sucrose in water solution measuring an osmolarity of 300 mOsm/kg were introduced in the chamber. The osmolarity was adjusted with osmometer (Gonotec Osmomat 3000 freezing point osmometer, Berlin, Germany) and filtered with Whatman PURADISC 30/0.2 CA s 0.2 μm filters. Electroformation then proceeded in three phases as described in (48), namely by a gradual increase of voltage from 0.02 V to 1.2 V over the course of 40 minutes at 10 Hz, followed by 90 minutes of constant electroformation at 1.2 V and 10 Hz, and ending in a vesicle-detachment at 1.3 V and 4 Hz for a minimum of 30 minutes. After electroswelling, the vesicle suspension, containing a final lipid concentration of 30 μM, was carefully collected with a pipette.

During in-bulk experiments, the GUV suspension was diluted 1:1 or 1:4 in an isotonic 300 mOsm/kg glucose solution. In experiments involving calcium, the glucose solutions contained 2, 5 or 10 mM CaCl_2_ while maintaining the value of the osmolarity fixed at 300 mOsm/kg.

#### 2.2.2 Multilamellar vesicles preparation for fluorescence spectroscopy

To prepare multilamellar vesicles (MLVs), 20 μL of 3 mM lipid in chloroform solution were spread at the bottom of a glass vial. The glass vial was then dried for 30 seconds under a nitrogen flow and placed in a vacuum pump for a minimum of an hour to evaporate the chloroform. After drying, 1 mL of 300 mOsm/kg sucrose was introduced in the vial. The vial was closed and left in a shaker for 30 minutes and then vortexed for 3 minutes to make MLVs. The MLVs were then diluted 1:1 with isoosmolar sucrose in cuvettes for observations. Fluorescence measurements were performed using a FluoroMax®-4 fluorometer (HORIBA Instruments Inc., USA).

### 2.3 Confocal microscopy

Fusion assays were carried out under a confocal microscope (Leica microsystems SP8 LIA Compact Supply Unit, Wetzlar, Germany) using a HC PL FLUOTAR 10×, NA 0.30 dry objective. To assess the permeability of the GUVs, we examined them under phase contrast using a HC PL FLUOTAR L 20x/0.40 dry objective. The vesicles fluorescently labeled with 0.5 mol % Atto488 DOPE were excited with solid state laser at 488 nm and the emission was detected in the 499-555 nm range with a PMT detector. Vesicles labeled with 0.5 mol% Atto633 DOPE were excited with a solid state laser at 638 nm and the emission was detected in a 649-714 nm range with a hybrid HyD detector. To minimize cross-talk for donor and acceptor excitation, the signal was collected in sequential mode. A third sequence was set up for the detection of FRET of the pair, exciting the dyes with the 488 nm laser and detecting within the Atto 633 DOPE emission range employing the same hybrid HyD detector. Simultaneously, transmitted light imaging was performed. Multiple xyt series were carried out at once using the Mark & Find function of the LAS X software of the microscope. Each measurement returned an output consisting of 4 channels (donor, acceptor, FRET and transmitted light), in which the signal was collected in the sequential mode.

### 2.4 Microfluidics set-up

Polydimethylsiloxane (PDMS) microfluidic chips were prepared according to a previously published protocol (66) by pouring a 10:1 mix of Sylgard™ Silicone Elastomer Base and Sylgard™ Silicone Elastomer Curing Agent (Biesterfeld Spezialchemie GmbH, Hamburg Germany) on a silicon wafer. The design of the microfluidic chip used was previously published (67). The silicone wafer engraved with this design includes 10 chips, each containing 12 channels with 16 rectangular traps each, which are made up of 19 posts respectively (see Fig. S1 in the supporting information, SI). As post shape and size may vary and be distorted in the process of etching the design on the silicon wafer, the post diameter measures 37 ± 0.5 μm, while the gap between the posts, standing at 6.5 ± 0.5 μm, is sufficiently small to trap vesicles with a diameter larger than 10 μm. The microfluidic devices were connected to a neMESYS Low Pressure Syringe Pump by CETONI GmbH, Korbussen Germany. The syringe was assembled with BASE 120 - neMESYS Basismodul pump by CETONI GmbH. To prevent GUVs from attaching to the glass slide, the PDMS chips were coated with BSA by loading 50-100 μL of a 2 mg/mL BSA in water solution in the chips to reduce nonspecific adhesion of GUVs to the glass slide and PDMS trap posts. The BSA was allowed to settle in the chip for 30 minutes, and then washed by flushing the volume of the chip (approximately 2 μl) with 300 mOsm/kg sucrose solution.

The GUVs were introduced into the microfluidic device by setting the flow rate to 5 μL/min. The microfluidic traps were considered sufficiently full if they displayed high local vesicle concentration, meaning that regions of the trap held GUVs in close proximity to one another. The loading stage was considered complete once we found a minimum of 8 sufficiently full traps (see examples in Fig. S1D-F of the SI), and the external glucose or calcium solutions were added to the inlet reservoir (Fig. S1A, B). From this step onward, the flow rate was reduced to 2 μL/min to continuously replace the external solution while reducing the mechanical forces acting on the trapped GUVs. If no solution replacement was ongoing, such as at the end of an experiment, the flowrate was maintained at 0.07 μL/min to prevent vesicle drifting. The flowrate was controlled using the neMESYS UserInterface software by CETONI GmbH. Observations were carried out by mounting the microfluidic chip on a confocal microscope. At these low flow rates, shear stress on the trapped GUVs is negligible and is not expected to influence membrane stability or fusion events.

### 2.5 Automated image analysis of vesicle fusion

We aimed to fuse vesicles of the same composition, specifically negatively charged GUVs. In each experiment, we measured the mean fluorescence emitted by vesicle clusters in separate traps of the microfluidic device over time. For image analysis of data collected on GUVs trapped in microfluidic chips, we employed a script designed to detect the boundaries of the regions occupied by GUVs and measure fluorescence intensity exclusively within those boundaries. This approach allowed us to track FRET evolution specifically within vesicle-containing areas rather than across the entire microfluidic chip, thereby reducing background noise and filtering out less relevant fluorescence signals. The mechanisms and details of this analysis can be found in section S2 of the SI and Fig. S2; we also apply it to bulk experiments (SI section S3 and Fig. S3). The script output consisted of three .csv files per trap, each containing the mean fluorescence intensity values for the fluorescence channels resulting from sequential scanning.

## 3 RESULTS AND DISCUSSION

As we aim to use FRET as an indicator of lipid mixing caused by calcium, we first identify a suitable FRET pair that minimizes false negatives due to spectral overlap. Specifically, we investigate the Atto488/Atto633 pair, comparing its emission and excitation spectra to the commonly used NBD/Rhodamine pair. After evaluating their spectra, we investigate how Atto488/Atto633-labeled GUVs appear under confocal microscopy when the dyes are mixed or separated, to verify their suitability for clearly reporting lipid mixing in calcium-exposed vesicles. In the subsequent sections, we use this FRET readout in microfluidic traps to quantify calcium-induced lipid mixing and observe early fusion events (Section 3.2). Next, we examine the role of negative-curvature lipids in promoting full fusion (Section 3.3), followed by the effects of membrane tension and cholesterol on fusion efficiency (Section 3.4). Together, these experiments provide a systematic assessment of how calcium, membrane composition, and mechanical factors govern the transition from lipid mixing to complete GUV fusion.

### 3.1 Atto488/Atto633 as FRET pair outperforms NBD/Rh eliminating donor emission contributions

Lipid mixing was monitored by measuring the FRET efficiency *E*_*FRET*_ (36):

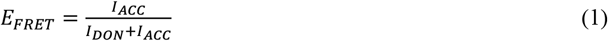

where *I*_*ACC*_ and *I*_*DON*_ are the intensities of the acceptor and the donor, respectively, when excited at the FRET donor’s excitation wavelength.

While NBD and rhodamine (Rh) are a typical and common FRET pair, we observed that vesicles containing only NBD exhibited significant fluorescence in the spectral range of Rh emission used to assess FRET. This strong overlap in the emission spectra introduced considerable fluorescence bleed-through even in vesicles containing exclusively NBD, as seen in Fig. 1B,C. To quantify the magnitude of this overlap, we performed fluorescence spectroscopy measurements on MLVs (Fig. 1A) and integrated the portion of NBD emission spectrum that falls in the emission range of Rh, i.e. the fraction of the donor emission collected in the FRET channel 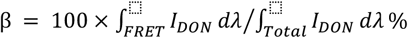 . Here, λ is the wavelength, the integral over the “FRET” range covers the wavelength range, in which the emission of the acceptor is collected in FRET measurements, while the integral over the “Total” range covers the whole wavelength range detected. The analysis was performed on the raw (not normalized) data shown in Fig. S4. For the NBD-Rh pair, we found β^*NBD*−*Rh*^ = 19.6%. This substantial spectral bleed-through means that even in the absence of any acceptor fluorophore, a strong signal may still be detected in the FRET channel, potentially leading to the erroneous interpretation of lipid mixing where none has occurred.

**Figure 1.**
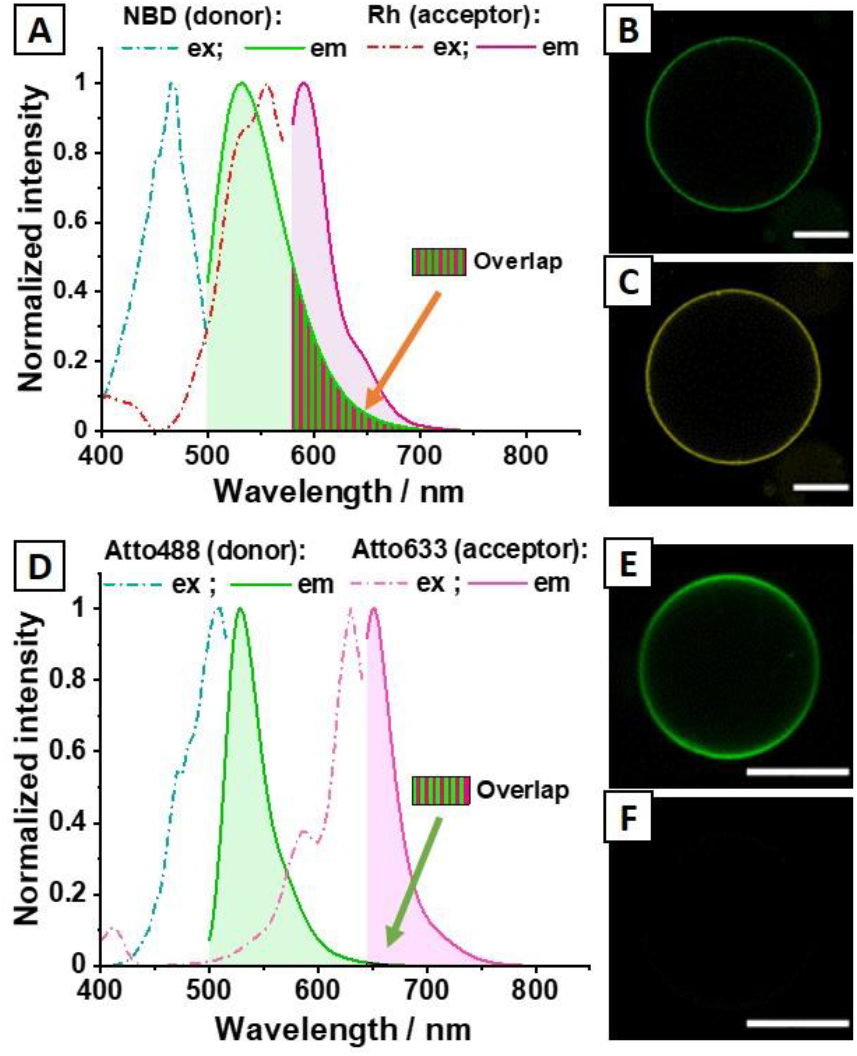
Atto488/Atto633 outperforms NBD/Rh as a FRET pair due to minimal spectral overlap. (A) Normalized fluorescence spectra of two MLV samples made of DOPC and containing 0.5 mol% of either NBD-DOPE (green) or Rh-DPPE (magenta); the dash-dotted and solid curves show the excitation and emission spectra, respectively. This FRET pair benefits from strong spectral overlap between the emissions of the donor (NBD) and the excitation of the acceptor (Rh). However, a significant portion of the emission signal of NBD falls in the collection range of the acceptor (Rh), as highlighted in green and red stripes. This overlap complicates the interpretation of the measured intensity in the FRET channel (assigned to *I*_*ACC*_ ), as it becomes unclear whether the signal originates from the donor or the acceptor. (B, C) Confocal microscopy images of a DOPC vesicle containing 0.5 mol% NBD-DOPE, imaged (B) in the NBD channel, with fluorescence signal collected between 498 and 537 nm, and (C) in the Rh channel, used to assess FRET signal, with collection between 562 and 620 nm. (D) Normalized fluorescence spectra of POPC MLV samples containing 0.5 mol% of either Atto488-DOPE (green) or Atto633-DOPE (magenta); the dash-dotted and solid curves show the excitation and emission spectra, respectively. This pair is characterized by well-separated and narrow spectral peaks, with minimal overlap between the emission spectra, as highlighted and hardly noticeable in the FRET detection range (649-714 nm). Importantly, while Atto488-DOPE emission overlaps with the excitation spectrum of Atto633-DOPE, there is no overlap between emission spectra of donor and acceptor. This ensures that the measured *I*_*ACC*_ is unambiguously emitted by the FRET acceptor, reducing signal contamination. (E, F) Confocal microscopy images of a 20:80 mol% POPS:POPC vesicle containing 0.5 mol% Atto488-DOPE, imaged in (E) the Atto488 channel, with fluorescence signal collected between 499 and 555 nm, and (F) the Atto633 channel, used to assess FRET signal, with collection between 649 and 714 nm. All scale bars: 20 μm. The raw data of the spectra in (A, D) are shown in Fig. S4.

We therefore explored the possibility of using Atto488-DOPE as a FRET donor and couple it with Atto633-DOPE as an acceptor. As shown in Fig. 1D, the emission of Atto488-DOPE overlaps with the excitation spectrum of Atto633-DOPE, permitting the use of these two fluorophores as a FRET pair. Furthermore, for the computed fraction of the emission spectra in the FRET channel (barely visible in Fig. 1F), we found β^*Atto*488−*Atto*633^ = 0.5%. Since only 0.5% of the Atto488-DOPE emission spectrum falls within the emission range of Atto633-DOPE, the overlap is very narrow, reducing ambiguity compared to the NBD-Rh FRET pair.

We also observed that Atto633-DOPE has a minor excitation peak under 500 nm (Fig. 1D), meaning that any vesicle containing Atto633-DOPE will emit some fluorescence when excited at shorter wavelengths, leading to signal detection in the FRET channel. Thus, to minimize direct excitation of the acceptor by the laser used for donor excitation, we selected an excitation wavelength of 638 nm for these experiments.

To demonstrate that Atto633 and Atto488 can serve as a FRET pair for microscopy, we prepared GUVs labeled with varying molar ratios of Atto488-DOPE and Atto633-DOPE (1:3, 1:2, 1:1, 2:1, 3:1), while maintaining the total amount of labeled lipids at 0.5 mol%. For comparison, and as a negative control (NC) we also prepared a mixture of GUVs labeled with only one of the two dyes, mimicking the pre-fusion state of individual vesicle populations. As shown in Fig. 2, GUVs labeled with both fluorophores consistently exhibit higher *E*_*FRET*_ values than the NC GUVs, regardless of fluorophore ratio. However, the *E*_*FRET*_ values of NC GUVs show considerable variability, with several outliers approaching the values of 2:1 or 1:1 Atto488:Atto 633 GUVs. This discrepancy arises from the mathematical definition of *E*_*FRET*_: since the donor fluorescence intensity (*I*_*DON*_) appears only in the denominator of Eq. 1, an imbalance where the number of donor-labeled GUVs is lower than that of acceptor-labeled GUVs can artificially increase the calculated *E*_*FRET*_ value. Note that this issue should also arise with LUV samples. The advantageous use of the Atto633 and Atto488 as a FRET pair is obvious when observing the GUVs under a confocal microscope: GUVs containing exclusively Atto488-DOPE or Atto633-DOPE show minimal or no signal in the FRET channel, whereas dual-labeled GUVs are clearly visible, see the inserts in Fig. 2. Analysis of the FRET signal was carried out using a script further explained in Fig. S3.

**Figure 2.**
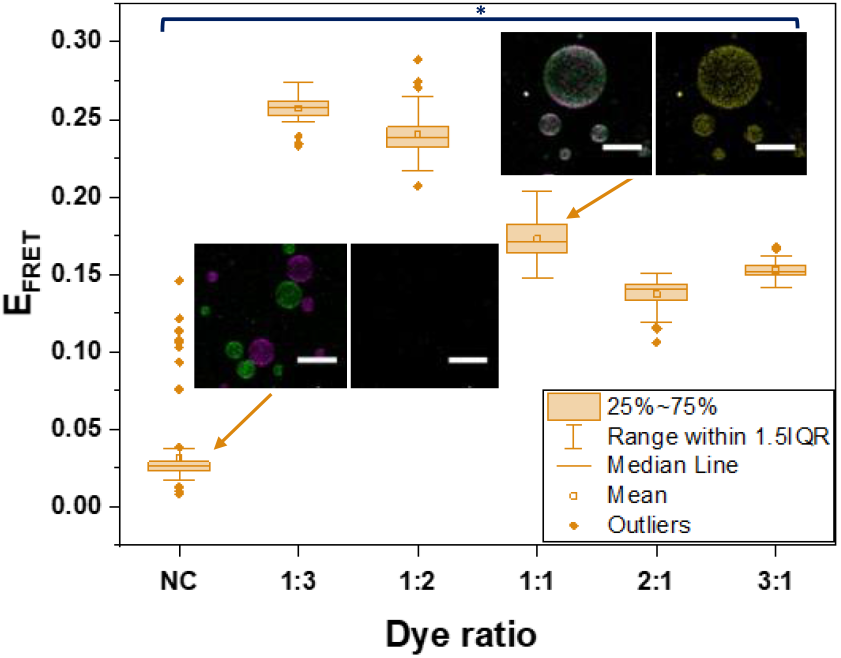
Validation of Atto488 and Atto633 as a FRET pair using dual-labeled GUVs. *E*_*FRET*_ measurement of GUVs containing various dye ratios show that when GUVs containing both Atto488 and Atto633 (at total amount of labeled lipids set to 0.5 mol% and varying molar ratios as indicated on the x-axis), FRET emission is considerably higher. NC acts as the negative control, corresponding to a mixture of GUVs labeled with only one of the two dyes. The lower left insert shows that the GUVs of the NC are visible in the Atto488 and Atto633 channels (merged image, top), but not in the FRET channel (bottom). Inversely, the upper right insert shows that GUVs containing both Atto633-DOPE and Atto488-DOPE each at 0.25 mol% are visible in the merged image (left) as well as in the FRET channel (right). Because of the proximity between the fluorophores present on the same vesicles, the GUVs are clearly visible in the FRET channel. Vesicles containing only one of the two fluorophores has returned markedly lower values than the GUVs holding a mix of both fluorophores. Scale bars 50 μm. On the average, 52 images per composition were measured. Box plots: central line shows the median; box represents Q1 (25th percentile) – Q3 (75th percentile); whiskers show lowest and highest values within 1.5× interquartile range of Q1/Q3; points beyond whiskers are outliers; squares show the mean value. Statistically relevant results are indicated by an asterisk * (Tukey test p<0.05). Analysis carried out on 306 GUVs in total.

Our results confirm that the spectral properties of Atto488 and Atto633 are well-suited for FRET-based detection of lipid mixing. As the fluorescent dyes are located on the lipid headgroup rather than on the acyl tails, the spectral overlap between the two dyes is sufficient to produce a measurable FRET signal upon close proximity (such as when lipid mixing occurs during membrane fusion) while their minimal emission cross-talk ensures that donor bleed-through into the FRET channel is negligible. This spectral separation significantly improves signal specificity, allowing us to attribute increases in FRET emission directly to lipid mixing rather than to optical artifacts. Based on these findings, we selected Atto488-DOPE and Atto633-DOPE as our FRET pair for all subsequent experiments. Employing microfluidic chips to confine and monitor clusters of GUVs over time, we then tracked the FRET signal variations following calcium addition and quantitatively assess the degree of lipid mixing during fusion.

### 3.2 Microfluidic-assisted analysis of calcium-induced lipid mixing in POPS/POPC GUVs reveals only limited fusion efficiency

To overcome the limitations of conventional single-vesicle assays, where random encounters between GUVs are rare and poorly controlled, we implemented a microfluidic trapping approach that enables controlled clustering of vesicles in defined geometries. This setup ensures that GUVs are in close proximity, an essential prerequisite for fusion, while allowing real-time observation of multiple vesicles under precisely defined conditions. In contrast to bulk assays or isolated GUV observations, where fusion events depend on stochastic collisions or uneven reagent distribution (also applying to LUV samples), the microfluidic platform facilitates reproducible initiation of fusion and high-throughput quantification across multiple vesicles simultaneously.

To investigate calcium-induced membrane interactions, we employed microfluidic chips containing PDMS posts that form traps (see Fig. S1). The traps enable prolonged imaging of vesicles while maintaining their position within the observation frame. This setup also allows full exchange of the GUV external solution, providing greater control over the vesicles’ environment. By confining GUV clusters in designated traps, we could monitor their FRET signal evolution over time under well-defined calcium exposure.

Figure 3A shows GUVs collected in one trap of the microfluidic device, and the boundaries within which fluorescence is measured; see also section S2 in the SI. Since *E*_*FRET*_ values can vary significantly between samples depending on the vesicle composition and arrangement within the trap, the absolute FRET signal is not a reliable standalone indicator. For instance, clusters with more donor- or acceptor-doped GUVs will display different baseline signals (see schematics in Fig. 3B and example images in Fig. S1D-F). To account for this variability, we analyzed the normalized change in FRET efficiency over time defined as 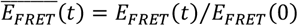, where *E*_*FRET*_(0) describes the signal in the trap upon initiating the GUV exposure to CaCl_2_. This relative metric provides a more robust comparison across traps and experiments, capturing the extent of lipid mixing independently of initial signal disparities.

**Figure 3.**
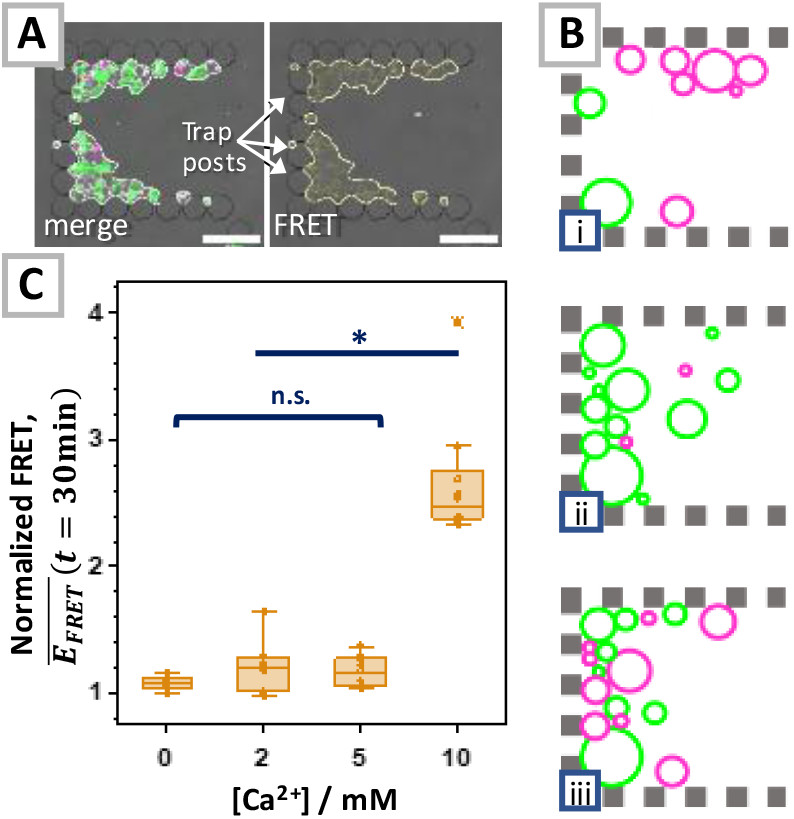
Microfluidic trapping enables quantitative analysis of calcium-induced lipid mixing in GUV clusters. (A) Confocal images of a microfluidic trap (overlaid bright field and fluorescence channels) containing a cluster of 20:80 POPS:POPS GUVs after 40 min exposure to 10 mM CaCl_2_. Bright-field imaging reveals the PDMS posts making up the trap. Left: composite fluorescence image showing donor (Atto488, green) and acceptor (Atto633, magenta) signals overplayed with the bright-filed signal. Right: merged image of the FRET (yellow) and bright-field signal. In both images, yellow outlines indicate the dynamic boundaries within which mean fluorescence intensity is measured over time (see also SI section S2); these boundaries adapt as vesicles move, fuse, burst, or escape. (B) Graphic representation of different initial conditions that determine the GUV cluster propensity for emitting FRET signal depending on the vesicle distribution in a microfluidic trap. (i) Either a nearly empty trap or one with segregated GUV populations, where donor and acceptor vesicles are unlikely to form contact points. (ii) A trap with a highly uneven donor-to-acceptor vesicle ratio, making lipid mixing difficult to detect. (iii) An optimal condition, where multiple contact points exist between vesicles of different populations, and donor and acceptor vesicles are evenly distributed. See Fig. S1D-F for example images illustrating the three cases. (C) Normalized FRET efficiency 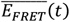 after 30 minutes of exposure to different calcium concentrations. Each point represents the mean value measured within one microfluidic trap. 29 GUV clusters measured. Box plot representation and statistical analysis as described in Fig. 2; in addition, statistically insignificant results are marked as “n.s.” (“non significant”, p>0.05). Data under the same bracket have all the same significance among each other, while data marked by the line shows the significance in relation between a group under a bracket or one set of data. In this case, the difference between 0, 2, and 5 mM CaCl_2_ are not significantly different from one another, while 10 mM is significantly different from 0, 2 or 5 mM. Analysis carried out on 306 GUVs in total.

We applied this approach to GUVs composed of 20:80 POPS:POPC, which represents a minimal model for negatively charged membranes. As shown in Fig. 3C, after 30 minutes of exposure to isoosmolar calcium solution of sufficiently high concentration, we observe a clear increase in normalized FRET. Although the FRET signal is highest at the contact regions between GUVs, we observe that FRET signal also increases at membrane regions exposed to the external solution and not in direct contact with other vesicles (see Fig. S5 in the SI), supporting lipid exchange between GUVs rather than mere adhesion. Notably, even with just 20 mol% POPS content, lipid mixing was detectable, indicating that a relatively low percentage of phosphatidylserine is sufficient to trigger fusion-like interactions between vesicles (see Fig. 3).

Despite normalization, the FRET signal shows significant spread across traps as reflected in the scatter of the data in Fig. 3C. This heterogeneity reflects the dynamic nature of the system: vesicles may shift, fuse, burst, or escape through trap gaps over time. Although an individual shift, fusion or bursting event is unlikely to significantly affect the overall value of 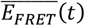 (see Fig. S6), their combination can affect the FRET readout. Crucially, however, an increase in FRET does not confirm full fusion - only lipid mixing.

Unlike conventional LUV-based fusion assays, where content mixing of encapsulated dyes is typically used to confirm fusion (36,40) but leakage can complicate interpretation, our microfluidic approach enables direct visualization of GUV morphology and topology to unambiguously identify fusion events. As shown in Fig. 3C, the normalized FRET signal increases at high calcium concentration, reaching values of 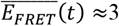, indicating lipid mixing. However, the corresponding confocal images show no evidence of full fusion: the vesicles maintain clearly defined boundaries, and only minor positional rearrangements are observed (see Movie S1 in the SI).

These observations suggest that although calcium promotes lipid mixing, it does not reliably promote full GUV-GUV fusion under the tested conditions. This underscores that calcium alone is a weak fusogen in this system. Given this, we shifted our focus to analysing additional factors that may influence the vesicle propensity to fuse, such as lipid composition and membrane tension, while recognizing that FRET-based measurements alone do not distinguish between lipid mixing and full fusion.

### 3.3 The role of negative-curvature lipids (PE) in calcium-induced vesicle fusion

Lipids with spontaneous negative curvature have been reported to facilitate membrane fusion (39), prompting us to investigate the impact of phosphatidylethanolamine (PE) lipids. Based on a prior report (48), we first tested 20:20:60 DOPS:DOPC:DOPE GUVs, here referred to as “test composition”. These GUVs are expected to predominantly burst and cluster upon calcium exposure, with occasional fusion events (48). In our observations, when subjected to 5 mM CaCl_2_ in a microfluidic chip, these GUVs underwent rapid and repeated fusion. However, this fusion cascade proceeded uncontrollably, with successive fusion events occurring in quick succession, ultimately leading to pervasive bursting that left the microfluidic trap empty and unusable (see Fig. 4 and Movie S2). Consistent with these observations, the normalized FRET traces for this composition (Fig. S7) rise well above the values measured for 20:80 POPS:POPC GUVs, reflecting the extensive lipid mixing that accompanies the rapid fusion cascade. Notably, the microscopy data indicate that full fusion events begin to appear when the normalized FRET signal 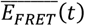 exceeds ∼3.5, providing an approximate threshold that aligns with the onset of visually confirmed fusion.

**Figure 4.**
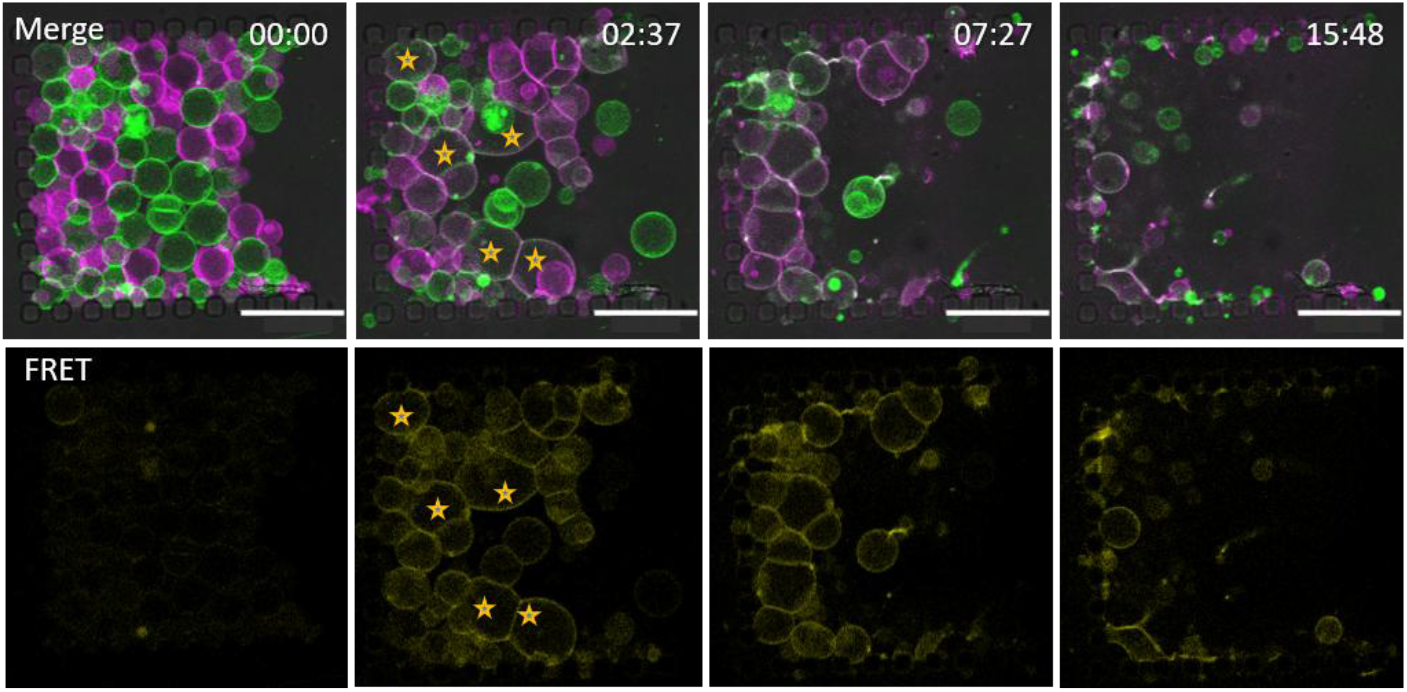
20:20:60 DOPS:DOPC:DOPE GUVs are fusogenic but mechanically unstable. The top images are overlay of bright-field and the two fluorescence channels (green and magenta) and the bottom images show the FRET signal; normalized FRET signal is given in Fig. S7, as well as vesicle cluster size. Upon exposure to 5 mM Ca^2+^ (second snapshot), the GUVs rapidly undergo a cascade of adhesion and fusion events, leading to the formation of large interconnected vesicles (some fused vesicles are marked with stars in the second snapshot). However, this extensive fusion is followed by vesicle bursting, ultimately leaving the microfluidic trap nearly empty; see Movie S2. Time stamps format: mm:ss. Scale bars: 100 μm.

Despite this composition’s lack of stability, we used it as a basis to further explore the role of negative curvature lipids in calcium-induced fusion. To maintain consistency with our previous experiments, we replaced DOPC and DOPS with their PO-lipid counterparts (POPC and POPS). These lipids, featuring one saturated and one unsaturated acyl chain, are widely used for GUV formation and are considered a more generalizable membrane model compared to fully saturated or fully unsaturated lipids, such as DPPC or DOPC. We opted not to apply this change to DOPE, as POPE has a transition temperature of 25 °C, which could affect membrane properties under our room-temperature experimental conditions.

Since PE lipids are typically enriched in the inner leaflet of biological membranes (32,68), they are not commonly used at high concentrations in GUV studies due to their limited physiological relevance in model systems. To address this, we explored compositions with reduced DOPE content as well. Figure 5 summarizes the observation from all explored systems.

**Figure 5.**
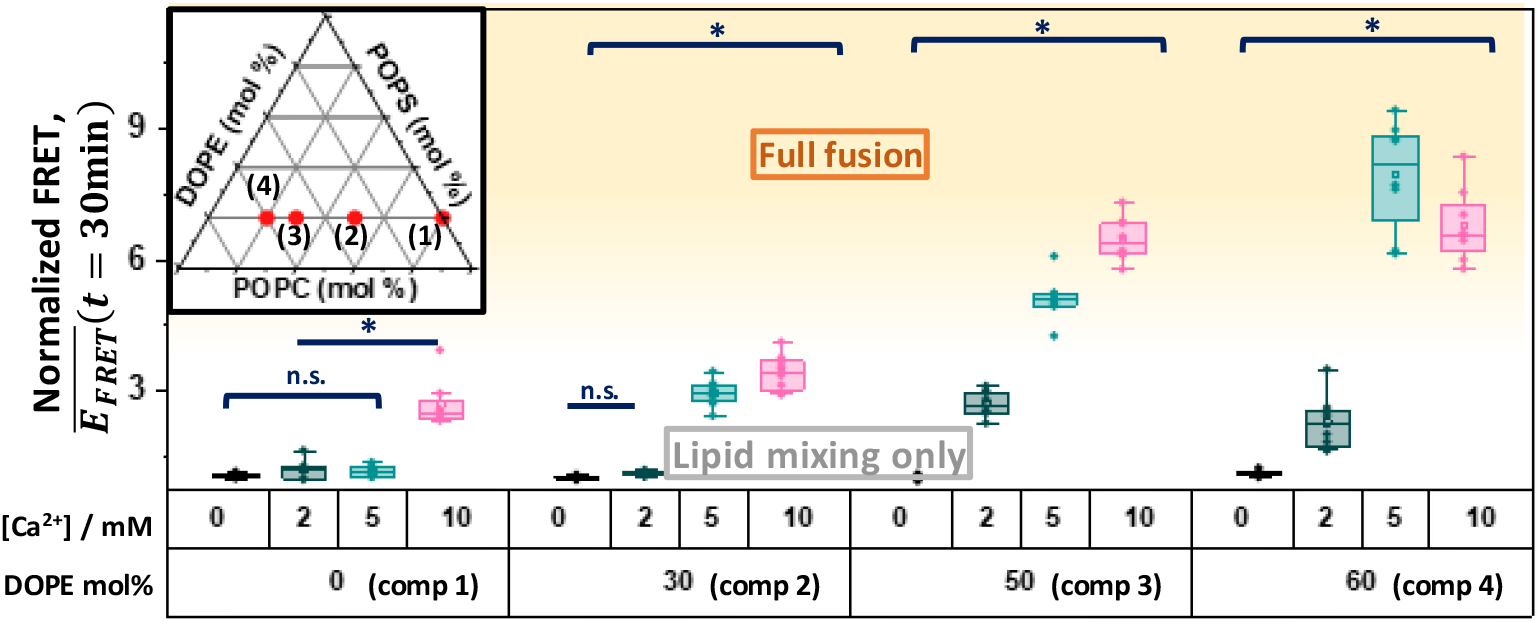
Full fusion requires high fractions of negative-curvature lipids. Influence of DOPE content and calcium concentration on lipid mixing and fusion in anionic GUVs composed of POPS:POPC:DOPE. The inset shows a ternary diagram with the four explored compositions (also listed in the table below the graph), used to assess how varying DOPE fractions influence the fusion propensity of anionic GUVs. The plot shows the normalized FRET signal measured after 30 min of exposure to different calcium concentrations. In all samples, POPS was kept constant at 20 mol%. The yellow-shaded region marks conditions under which full fusion events became increasingly frequent as directly observed in the microscopy sequences, whereas only lipid mixing occurred below this threshold. Box plot representation and statistical analysis as described in Figs. 2 and 3. Analysis carried out on 125 GUV clusters in total.

As shown in Fig. 5, GUVs containing 30 mol% DOPE displayed no notable increase in fusion propensity (see example in Fig. S8) compared to the previous measurements (Fig. 3), and no fusion events were observed over the recorded time series. Under the microscope, these vesicles remained stable across all tested calcium concentrations. The results indicate that the presence of 30 mol% DOPE does not significantly enhance calcium-induced fusion.

Conversely, GUVs containing 50–60 mol% DOPE exhibited markedly increased sensitivity to calcium. Even at 2 mM CaCl_2_, substantial lipid mixing was detectable, and at 5 or 10 mM CaCl_2_, the mean normalized FRET signal was, on average, fourfold higher than in DOPE-free GUVs, reaching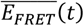 values characteristic for fusion. Consistent with this, time-series recordings revealed frequent fusion events at calcium concentrations of 5 mM and above. Unlike the test composition (48), these GUVs did not undergo rapid and pervasive bursting after fusion initiation. Instead, most vesicles remained intact post-fusion. However, phase-contrast imaging revealed partial leakage in some GUVs, indicated by loss of optical contrast. This contrast loss likely reflects disruption of the initial sugar asymmetry: intact vesicles filled with sucrose appear dark with a bright rim, whereas leaky vesicles lose this characteristic halo as the internal and external solutions equilibrate. An example is provided in Fig. 6 and Movie S3, with additional cases provided in Fig. S9.

**Figure 6.**
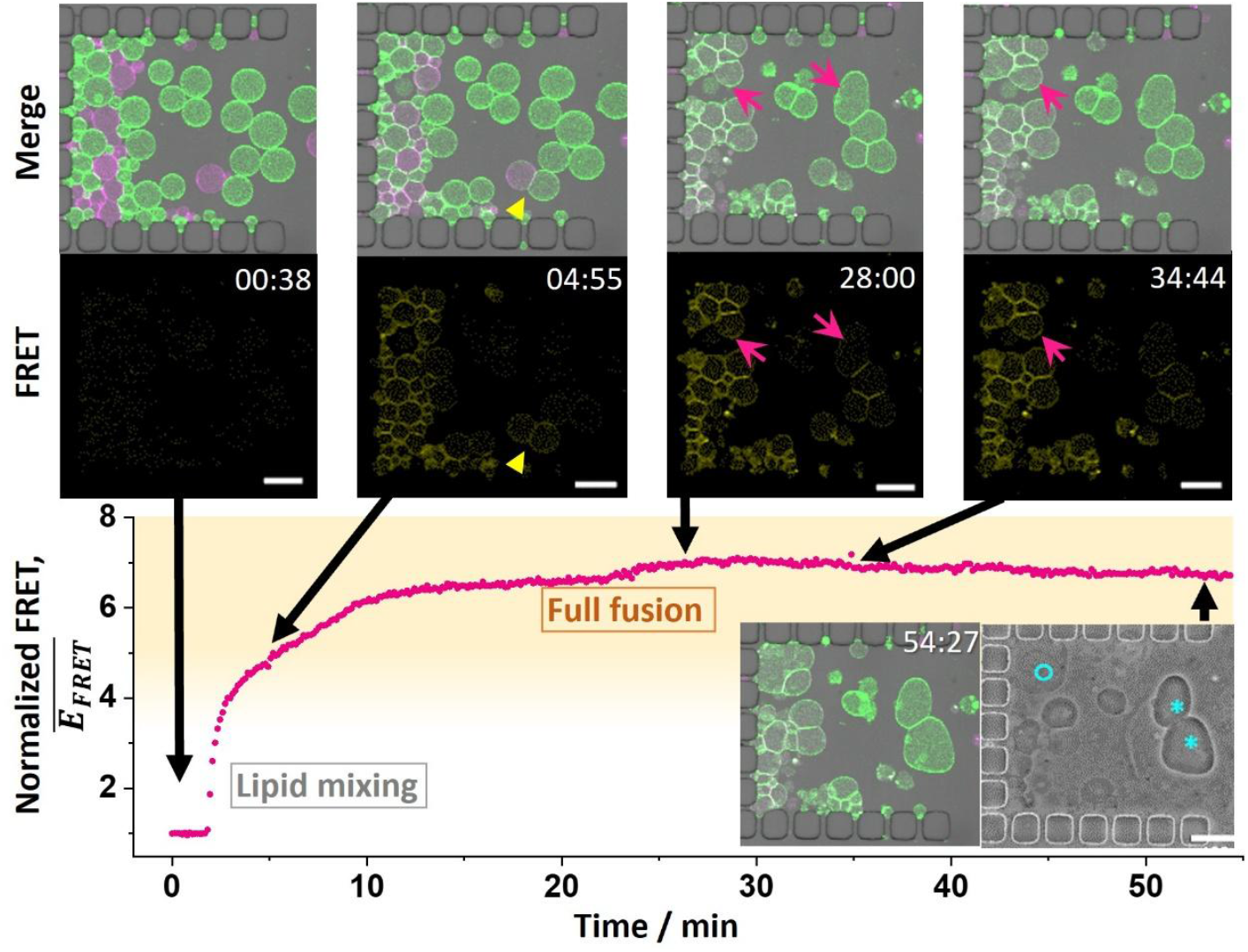
Balancing negative curvature lipids and chain saturation enables stable full fusion without GUV rupture. The graph shows progressive increase in lipid mixing, with representative insets illustrating the fusion cascade. Top row of images shows Atto488 (green) and Atto633 (magenta) merged fluorescence and bright field signal, bottom – FRET channel overlaid with bright field signal. The magenta arrowheads mark individual fusion events. A yellow arrowhead highlights a vesicle pair in which FRET signal is clearly visible beyond the contact interface, indicating lipid redistribution over the entire vesicle membrane rather than confinement to the docking region. The graphs shows the evolution of the normalized FRET signal following the addition of 10 mM CalCl_2_ to 20:20:60 POPS:POPC:DOPE GUVs. In contrast to DOPS:DOPC:DOPE GUVs (Fig. 4), these POPS:POPC-containing vesicles remain largely intact after fusion. Even 30 min after the onset of fusion, numerous fused GUVs persist within the trap, with minimal bursting. The accompanying phase-contrast image recorded at t = 55 min (inset, right) confirms that sugar asymmetry was preserved in some vesicles (asterisks), indicating they remained sealed, while others showed leakage (empty cyan circle). This sequence is also shown in Movie S3; additional examples at different conditions are shown in Fig. S9. Time stamps format: mm:ss Scale bars: 50 μm.

We find that fusion events can typically be observed when the normalized FRET signal 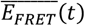 exceeds ∼3.5. Although this value alone does not define fusion, it correlates with GUVs that have undergone full fusion as opposed to lipid exchange through hemifusion (see Fig. S10). Thus, 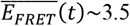 serves as a qualitative indicator that requires verification through direct observation of confocal images, rather than a parameter that determines whether fusion has occurred.

The combined results of in Figs. 5 and 6 indicate that while DOPE facilitates calcium-induced fusion, its effectiveness is strongly composition-dependent, requiring both high negative curvature lipids and sufficient stability to prevent bursting without blocking full fusion. Notably, GUVs composed of POPS:POPC:DOPE demonstrated greater stability than their DOPS:DOPC:DOPE counterparts (the test composition). Even after one hour of observation of the former composition, the microfluidic traps retained a significant number of fused vesicles (Figs. 6, S9 and S11). Unlike the rapid and uncontrollable bursting seen in the test composition (Fig. 4), these vesicles exhibited limited bursting and, in some cases, maintained their structural integrity after undergoing full fusion, as confirmed by phase-contrast imaging.

### 3.4 Membrane tension and cholesterol modulate fusion efficiency

We next examined how membrane tension and cholesterol content influence fusion, using our microfluidic trapping and analysis platform. Membrane tension is widely recognized as a key promoter of membrane fusion (69,70). Because several of our conditions produced lipid mixing without progressing to full fusion (see Fig. 5 and Movie S1), we tested whether increasing membrane tension would drive the transition from hemifusion to complete fusion by triggering the opening of the fusion pore. Because elevated membrane tension has been reported to suppress hemifusion and reduce lipid mixing efficiency (71), we deliberately introduced tension only after lipid mixing had been established, in order to specifically test its effect on the hemifusion-to-fusion transition.

To increase tension, we subjected GUVs to a hypoosmolar environment in the presence of calcium, leading to vesicle inflation and increased membrane tension. We selected GUV compositions and calcium concentrations where lipid mixing occurred but without full fusion: 20:20:60 POPS:POPC:DOPE at 2 mM CaCl_2_, and 20:50:30 POPS:POPC:DOPE at 2 mM and 5 mM CaCl_2_. The trapped GUVs were first exposed to an isoosmolar calcium-containing solution (303 mOsm/kg). To increase membrane tension, we then introduced a hypotonic solution (257 mOsm/kg) while keeping the calcium concentration constant. This osmotic shift increases GUV volume by ∼16%, corresponding to an estimated ∼10% increase in vesicle area. Assuming an initially spherical GUV with no excess membrane reservoirs (e.g. stored in nanotubes or defects), such stretching would elevate the membrane tension substantially and is likely to induce bursting. In practice, electroformed anionic GUVs may contain excess membrane area in the form of sub-resolution nanotubes (72), which would partially buffer the imposed osmotic strain. In this case, the estimated area increase represents an upper bound for membrane stretching. Nevertheless, the observation of vesicle rupture during inflation (Movie S4) indicates that a significant fraction of vesicles reached tensions close to the lysis threshold, confirming that the applied osmotic gradient generates substantial membrane tension.

Using typical stretching elasticity moduli of 130–200 mN/m, with 200 mN/m being typical for pure POPC membranes (73) and 130 mN/m for POPC membranes containing 20 mol% charged lipids (74), the imposed osmotic stress translates to a tension increase of ∼13–20 mN/m, sufficient to reach or exceed the lysis limit. Consistent with this estimate, several vesicles ruptured during inflation (see Movie S4). Despite the considerable stress, GUVs that did not burst remained strongly adhered to neighboring vesicles rather than detaching to regain a more energetically favorable spherical shape (see Fig. S12), demonstrating the stability of the docking interfaces under experimental conditions.

FRET efficiency measurements remained consistent with previous experiments, collectively indicating that higher membrane tension enhances fusion propensity—but only when the membrane contains a high fraction of DOPE (see Fig. S12). Notably, GUVs containing 60 mol% DOPE exhibited increased propensity to fuse under hypoosmotic 2 mM [Ca^2+^], whereas vesicles with lower DOPE content did not.

We then turned to cholesterol as a key regulator of membrane mechanics. Cholesterol is abundant in eukaryotic membranes and modulates both fluidity and bending rigidity, yet its role in membrane fusion remains controversial. It can either promote fusion by inducing negative curvature in certain lipid environments or suppress it by increasing packing and stiffness, see e.g. (75). To explore this, we replaced DOPE with cholesterol in our model composition, forming 20:50:30 POPS:POPC:Chol GUVs. While these vesicles exhibited lipid mixing similar to that of membranes containing ≤50 mol% DOPE (see Fig. S12), no full fusion events were observed.

Together, these results highlight that while increasing membrane tension can facilitate fusion in PE-rich membranes, cholesterol-rich membranes resist progression to full fusion. Although lipid mixing still occurs, suggesting that early intermediates such as stalk formation are accessible, cholesterol likely stabilizes hemifusion-like states and increases the energetic barrier for fusion pore nucleation and expansion (76). By increasing membrane order, bending rigidity, and resistance to topological rearrangements, cholesterol appears to suppress the final transition to full fusion. This interplay between mechanical stress and molecular composition emphasizes the delicate balance that has to be established in synthetic systems to control fusion efficiency.

## 4 DISCUSSION AND CONCLUSION

In this study, we introduce a microfluidics-based approach to investigate calcium-induced fusion of anionic GUVs. The use of microfluidic chips offers several key advantages: they enable long-term observation of individual vesicles under stable conditions, allow precise control over environmental conditions through complete and rapid solution exchange, and facilitate vesicle confinement that promotes clustering, an essential precursor of fusion. In contrast, bulk assays using LUV suspensions or single-GUV experiments relying on local solution injection often suffer from poorly defined conditions, including concentration gradients of vesicles or fusogen concentrations, inhomogeneous diffusion of external solutions, and uncontrolled flow artifacts. Additionally, bulk content-mixing assays can generate misleading results, as transient vesicle bursting and reclosure events, which are undetectable in ensemble measurements, may be erroneously interpreted as fusion (40). By confining GUVs in microfluidic traps, our platform enables an unambiguous distinction between lipid mixing and full fusion across large vesicle populations, while preserving membrane integrity and allowing direct visualization of dynamic fusion events.

A key strength of our combined approach is the ability to cross-validate FRET-based lipid-mixing measurements with direct microscopic evidence of membrane merger. The dual approach - combining quantitative FRET measurements with direct topological observation, strengthens the reliability of fusion analysis and overcomes the limitations of FRET-only assays. While FRET provides a sensitive and high-throughput readout, it inherently overestimates fusion efficiency because it only reports mixing between differentially labeled vesicles; fusion between vesicles carrying identical probes remains invisible (the green-labeled vesicles fusing on Fig. 6, snapshot at 28 min demonstrates a fusion event undetectable by FRET); this is a limitation shared with all bulk LUV studies. By correlating FRET data with microscopy, we identified an empirical threshold above which full fusion becomes detectable by imaging: normalized FRET values of 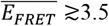 consistently correspond to complete fusion events. This empirical benchmark can be adopted in future studies, where microscopy-validated FRET thresholds could serve as a generalizable calibration tool for distinguishing lipid mixing from full fusion in population-level assays.

The exploitation of direct microscopic observations could be further improved by labeling the GUV aqueous contents to more accurately monitor vesicle number and size. However, this approach is limited to vesicles that remain impermeable throughout the experiment. Moreover, confocal imaging in a single plane typically captures only a cross-section of the vesicle, which may not correspond to the equatorial section, and vesicle adhesion and shape changes during hemifusion or after fusion further complicate size estimation. In principle, 3D imaging approaches, such as dynamic light-sheet microscopy, could provide more precise measurements of vesicle volume and fusion dynamics, though integration with microfluidic trapping remains to be explored.

Using this integrated strategy, we showed that full fusion occurs reliably only in GUVs containing ≥50 mol% DOPE (with POPS and POPC) at specific calcium concentrations (Figs. 5, 6, and S9). Unlike the behavior observed in membranes composed entirely of unsaturated lipids (Fig. 4), fused PE-rich vesicles largely remained intact, with only sporadic leakage detectable via changes in phase contrast (Figs. 6 and S9). This result highlights the critical role of PE lipids (characterized by strong negative spontaneous curvature) in promoting fusion, aligning with previous findings in LUV studies (27). Notably, PC lipids strongly inhibit calcium-induced fusion, even when present in small amounts in anionic LUVs (27), an inhibitory effect intensified in GUVs owing to their lower inherent curvature. Conversely, LUVs, by virtue of their small size and high curvature, presumably require less PE to stabilize the curved intermediates of fusion. Thus, both curvature and membrane composition strongly modulate the fusogenicity of calcium, making GUVs a more stringent and physiologically relevant platform for exploring protein-free fusion pathways.

While calcium alone is insufficient to drive full fusion in anionic GUVs, our data reveal that it robustly promotes early fusion intermediates such as vesicle clustering and extensive lipid mixing. These early-stage effects appear broadly independent of composition at ≥5 mM Ca^2+^ (Figs. 3, 5, S8, S9). We further examined whether membrane tension could serve as a secondary trigger to complete fusion, given that tension is widely recognized to lower energy barriers along the fusion pathway. However, even substantial tension increases induced by hypoosmotic inflation were unable to induce full fusion unless the membranes already contained large amounts of DOPE. This highlights the dominant influence of negative-curvature lipids over tension in determining fusion outcomes.

Beyond clarifying the fusogenic action of calcium, our work also emphasizes important limitations of commonly used FRET-based fusion assays. We showed that the conventional NBD/Rh pair is not ideal and exhibits spectral bleed-through (Fig. 1). Then we showed that a significant fraction of the detected FRET signal may arise from lipid mixing and rupture rather than true topological merger. GUVs, with their optical accessibility, offer a unique advantage here: fusion, hemifusion, rupture, and reclosure events can be directly discriminated, helping reinterpret observations from LUV-based assays that lack topological resolution. In this context, GUVs can serve as a “blown-up” version of LUVs, offering a visual window into processes otherwise masked in bulk. However, it should be noted that the dynamics of GUV-GUV fusion cannot be directly compared to LUV-based assays. LUV experiments are typically performed at lipid concentrations two orders of magnitude higher than in GUV systems, resulting in frequent vesicle encounters and different lipid-to-calcium ratios. In contrast, GUVs are spatially separated and practically immobile in traps, and fusion can only occur when vesicles are in contact. Furthermore, calcium delivery in microfluidic traps is not instantaneous and may reach different clusters at slightly different times, producing variability in early fusion kinetics. Ensemble measurements of LUVs also cannot account for vesicle rupture or population loss over time, which can affect the observed FRET signal. Consequently, while the timing of fusion events can be reported for GUVs, these kinetics are highly situation-dependent and cannot be directly compared to LUV assays (Fig. S11).

Overall, our findings indicate that calcium alone is insufficient to drive full fusion in anionic GUVs unless the membrane composition is finely tuned to balance charge, stability, and negative curvature lipids. While calcium remains a viable protein-free fusogen, its effectiveness is far more limited in GUVs than previously assumed. Our results call for a reassessment of assumptions derived from LUV studies and show that fusion in GUVs requires a delicate interplay of multiple factors, rather than a simple reliance on calcium as a universal fusogen.

Membrane fusion plays a foundational role in synthetic biology, particularly in efforts aimed at constructing growing, dividing, or communicating minimal cells. Synthetic cells require mechanisms for controlled, repeated, and non-destructive membrane growth, yet most available fusion triggers either fail to yield sustained fusion or compromise compartment integrity. Our results provide quantitative guidance for implementing calcium-based fusion pathways in bottom-up cell systems. Specifically, the ability to tune fusion efficiency by adjusting PE content and calcium concentration, combined with the microscopy-validated FRET threshold (∼3.5) for detecting full fusion, offers a practical framework for engineering fusion-driven membrane expansion in a reproducible manner. More broadly, the microfluidic platform developed here enables systematic exploration of fusogens, membrane compositions, and environmental cues, advancing the development of synthetic cells capable of controlled membrane remodeling, nutrient uptake, and growth.

## 5 AUTHOR CONTRIBUTIONS

RD proposed and supervised the project. MA and RD designed the experiments. MA and RH performed the experiments. MA analyzed the data. NY and SP provided instruction and direction on the use of microfluidic devices. MA and RD wrote the manuscript. All authors read and agreed with the final version of the manuscript.

## 7 ACKNOWLEDGEMENTS

This work relates to Department of Navy award N62909-22-1-2027 issued by the Office of Naval Research.

