## Supporting Information for "Protein-free membrane fusion: a refined view of the delicate fusogenic properties of calcium"

### S1. MICROFLUIDIC DEVICES

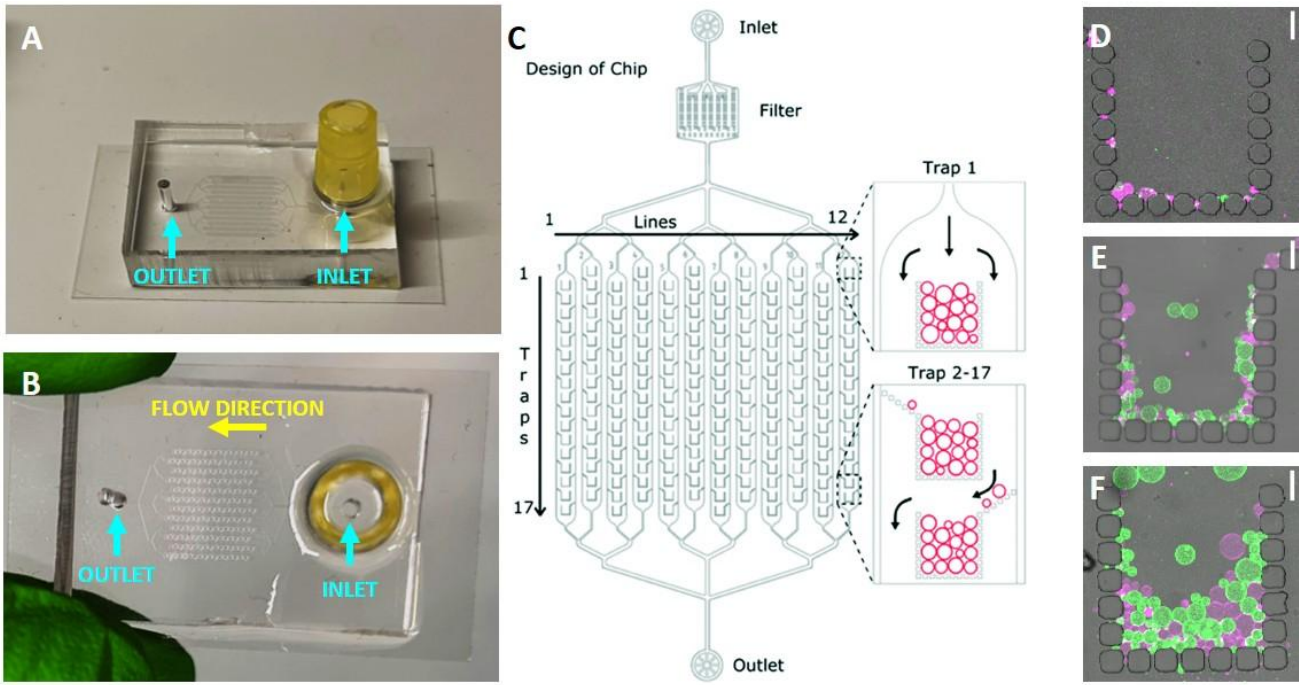


**Figure S1. Microfluidic chips: design and loading.** (A, B) Side and top views of the microfluidic chip setup. A reservoir is formed by cutting off the bottom of a plastic pipette tip (yellow) and attaching it onto the chip inlet using uncured PDMS. For surface treatment, a 2 mg/mL BSA solution in water is added to the reservoir, and the chip is centrifuged using a Rotina 420R centrifuge, produced by Unity™ Lab Services by Thermo Fisher Scientific, Waltam Massachussets, US. The centrifuge is set at 900 rpm for 10 minutes at 23 °C; ramp-up and ramp-down parameters both set at 9. (C) Sketch of the original microfluidic chip design, adapted from (1) with permission from the Royal Society of Chemistry. In this study, a modified version was used, featuring 12 channels, each with 16 rectangular traps composed of 19 posts. The posts have dimensions of 35 µm in diameter and a gap distance of approximately 5 µm. (D–F) Representative examples (overlaid bright-field and fluorescence images of two differently labeled GUV populations) of trap loading quality prior to calcium exposure. (D) An underloaded trap with minimal vesicle–vesicle contact; unsuitable for analysis. Shown: 20:80 POPS:POPC GUVs. (E) A moderately loaded trap with sufficient GUV contact points to allow data collection if better traps are unavailable. Shown: 20:20:60 POPS:POPC:DOPE GUVs. (F) An optimally loaded trap, with numerous GUV–GUV contacts suitable for robust measurements. Shown: 20:50:30 POPS:POPC:DOPE GUVs. In all of the examples, a fraction of the vesicles is labeled with 0.5 mol % of either Atto 488 DOPE (donor, green) and Atto 633 DOPE (acceptor, magenta). Scale bars: 50 µm.

### S2. MARK-AND-FIND SEQUENCE PROCESSING FOR MICROFLUIDICS EXPERIMENTS

To uniform the image analysis process, we wrote a program with the built-in ImageJ scripting tool. The commented code was uploaded on OwnCloud and can be found at https://owncloud.gwdg.de/index.php/s/QRqGxX4cDIPuncp.

To identify the trap regions in which GUVs are present, the donor (imaging channel, CH00) and acceptor (CH02) contrasts are increased, making both donor and acceptor populations distinguishable. The images are added so that the program will identify the presence of both GUV types indiscriminately. To assess the general area of the GUVs, the image is then put under a band-pass filter to blur the lines, and then binarized. We made use of ImageJ "Analyze>Analyze particles…" function to create a region of interest (ROI) selection. See Fig. S2 for a visual representation of this process.

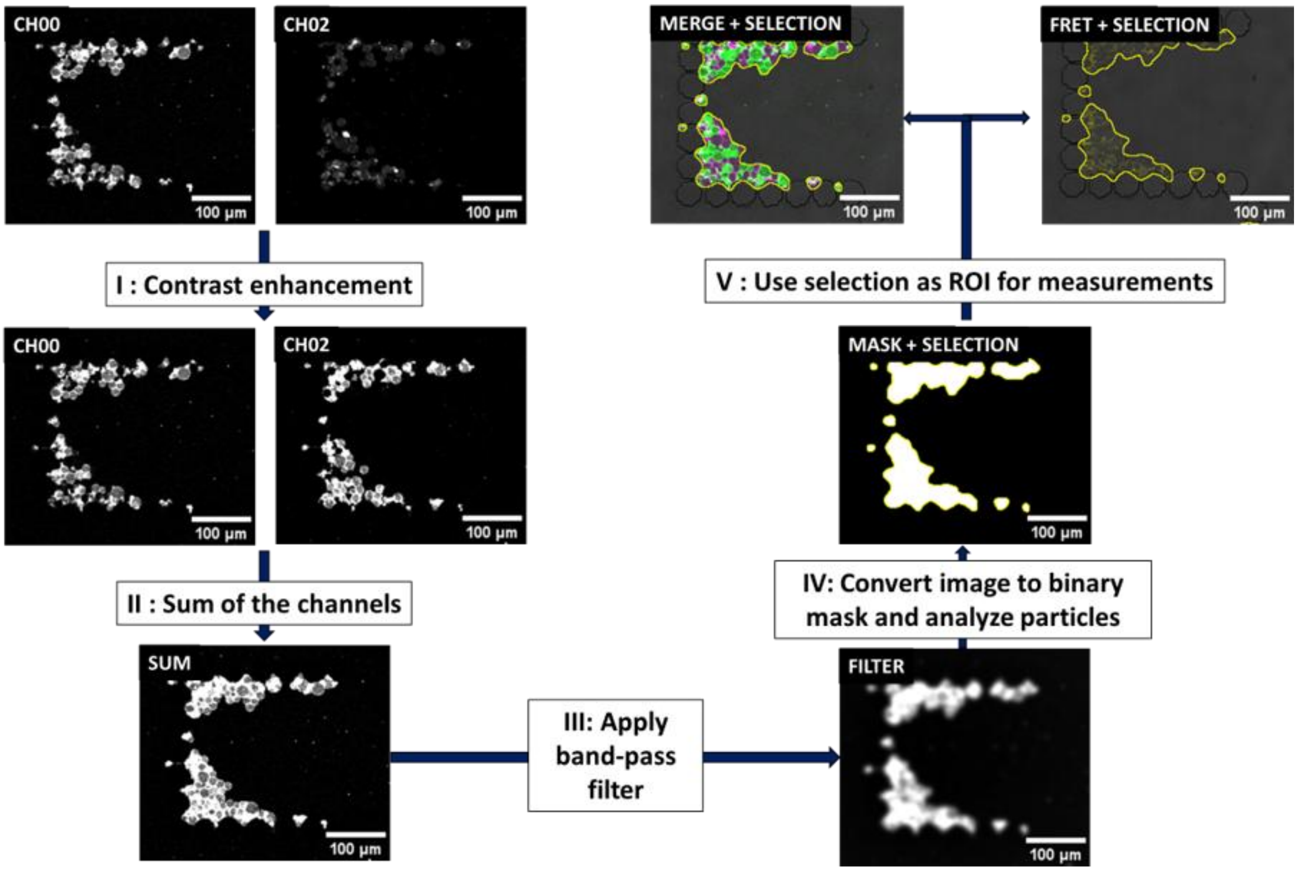


**Figure S2. Workflow for automated ROI selection and fluorescence quantification in GUV imaging in microfluidic chips.** Image analysis pipeline used by the custom ImageJ script to quantify fluorescence signals from GUVs while minimizing background noise. (I) Each fluorescence channel is contrast-enhanced individually. (II) The enhanced images are summed to produce a composite image that highlights all GUVs, independent of fluorophore identity. (III) A band-pass filter is applied to soften vesicle edges, aiding in consistent object recognition. (IV) The resulting image is binarized, and ImageJ's "Analyze Particles…" function is used to automatically define ROIs encompassing only the GUVs. (V) These ROIs are then used to extract fluorescence signal intensities within vesicle boundaries for subsequent analysis.

### S3. TILE-SCAN BASED PROCESSING FOR IN-BULK EXPERIMENTS

The processing script to determine the ROI in bulk experiments functions with the same principle as displayed in Fig. S2, Fig. S3 provides a graphic depiction of the process.

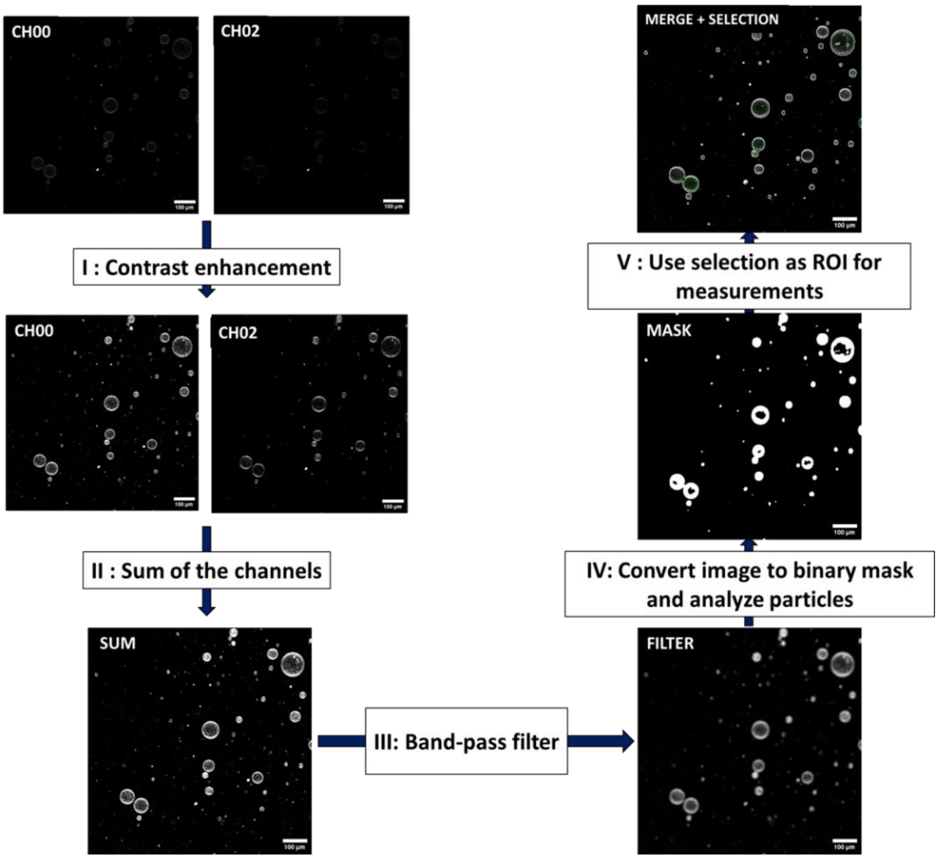


**Figure S3. Image analysis workflow for ROI-based fluorescence quantification of GUVs in bulk experiments.** Illustration of the image analysis procedure implemented in the ImageJ script to quantify fluorescence signals from GUVs. To minimize background noise, fluorescence is measured only within regions of interest (ROIs) that tightly surround the vesicles. These ROIs are automatically generated based on contrast-enhancing and filtering steps (similar to those in Fig. S2) that ensure consistent and accurate signal extraction from individual GUVs across fluorescence channels.

### S4. SPECTRAL ANALYSIS OF ATTO488/ATTO633 FRET PAIR

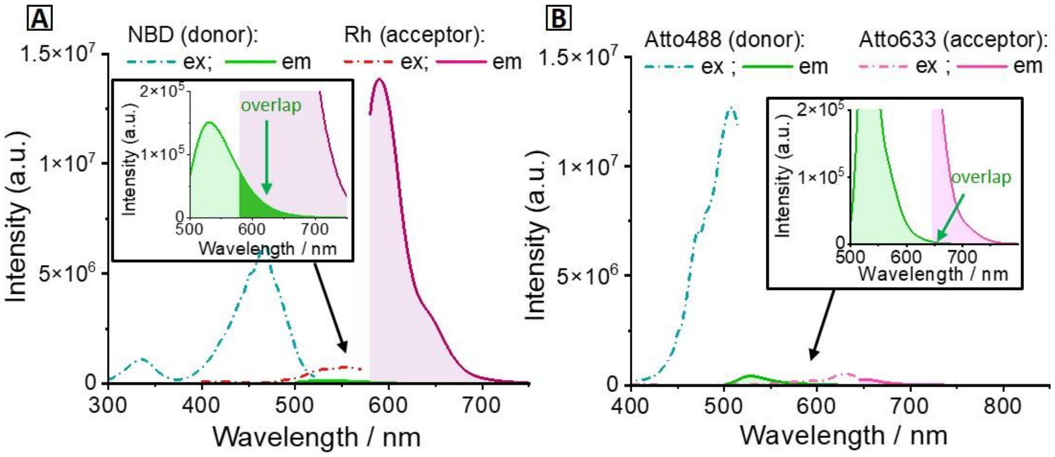


**Figure S4. Raw fluorescence spectra of the two FRET pairs NBD/Rh and Atto488/Atto633.** The corresponding normalized spectra are shown in Fig. 1 in the main text. (A) Excitation (dash-dotted) and emission (solid) spectra of DOPC MLVs containing either 0.5 mol% NBD-DOPE (green) or Rh-DPPE (magenta). A substantial fraction of the NBD emission falls within the wavelength range used to collect the Rh acceptor signal, as indicated by the dark green highlighted region in the inset. (B) Excitation (dash-dotted) and emission (solid) spectra of POPC MLVs containing either 0.5 mol% Atto488-DOPE (green) or Atto633-DOPE (magenta). In contrast to the NBD/Rh pair, spectral bleed-through is negligible, as shown in the inset.

### S5. DISTINGUISHING ADHESION AND LIPID MIXING

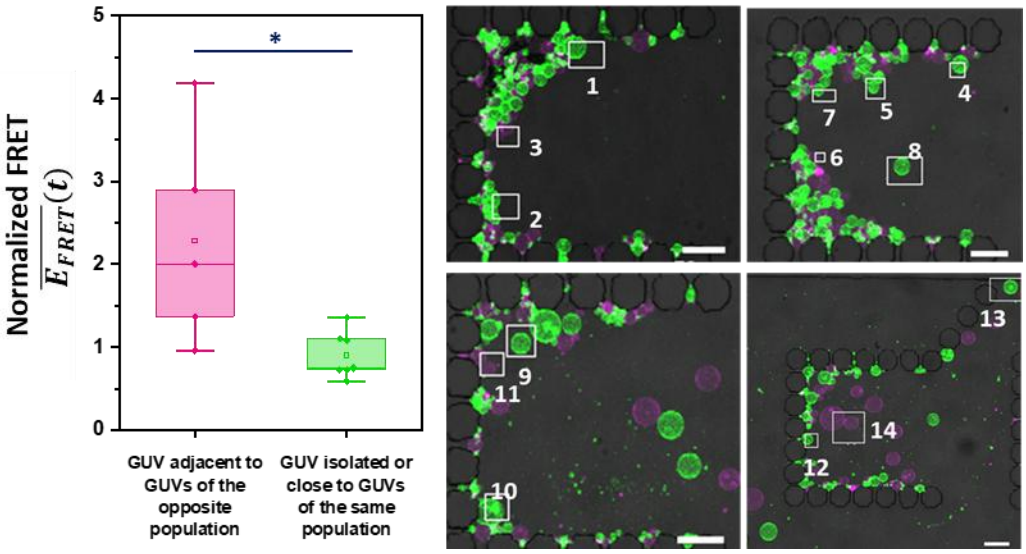


**Figure S5. GUVs (20:80 POPS:POPC ) adjacent to GUVs of a different population display higher values than GUVs that are either isolated or in contact only with GUVs of the same type even when excluding interpopulation adhesion.** To determine whether the increase in FRET signal in microfluidic traps can be exclusively attributed to the areas where GUVs of different populations are in contact with one another, we segmented our recordings and measured the normalized FRET signal $\overline{E_{FRET}}$ within limited areas, intentionally excluding the areas in which Atto488/Atto633 GUV pairings are in contact, as shown on the right, where the segmented areas have been outlined in white. One group of segments measures the FRET signal of vesicles that come into contact with at least one neighbor vesicle containing a different fluorophore (e.g. segments 4, 6, 7; pink bar in the graph), while the other category includes GUVs that are either isolated or are surrounded by GUVs with the same label (e.g. 5, 8, 14; green bar). We see that, despite excluding Atto488/Atto633 contact areas, GUVs with interpopulation contact interfaces exhibit a higher $\overline{E_{FRET}}(t)$ value than GUVs that are isolated or with same-labelled GUVs. This shows that the GUVs that are attached to other GUVs of other labels exhibit an increased FRET signal that cannot be attributed to adhesion or to an unforeseen effect of calcium indiscriminately affecting the fluorescent labels, but rather to the exchange of lipids between GUVs following $Ca^{2+}$ addition. 12 segments analyzed. Box plot representation and statistical analysis as described in Fig. 2. Scale bars: 50 µm.

### S6. EFFECT OF SINGLE BURSTING EVENTS ON OVERALL FRET SIGNAL

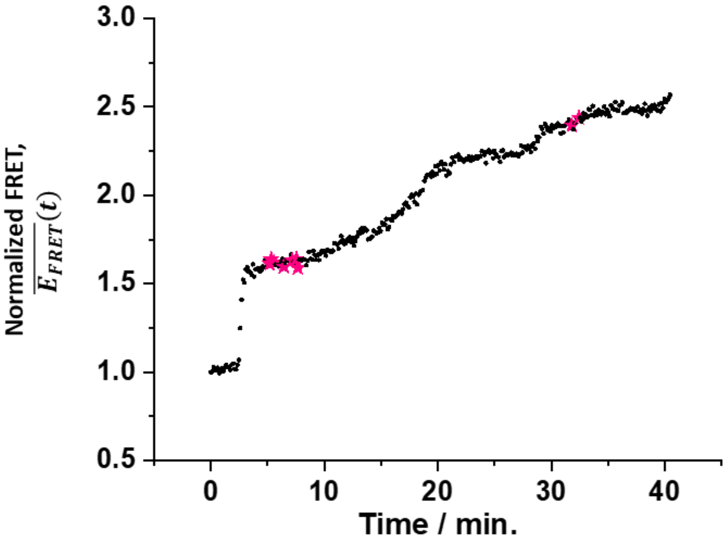


**Figure S6. Single bursting events have limited impact on the normalized FRET signal $\overline{E_{FRET}}(t)$.** Time evolution of $\overline{E_{FRET}}(t)$ measured in a microfluidic trap containing 20:80 POPS:POPC GUVs after addition of 10 mM $Ca^{2+}$. Time points immediately following visually confirmed bursting events are marked with magenta stars. While such events induce transient fluctuations in the FRET signal, their effect remains limited because $\overline{E_{FRET}}(t)$ is calculated from mean fluorescence intensities across the entire trap. Sustained increases in $\overline{E_{FRET}}(t)$ are therefore unlikely to arise from isolated rupture events alone.

### S7. NORMALIZED FRET INCREASE IN BURSTING-PRONE GUVs DURING FUSION

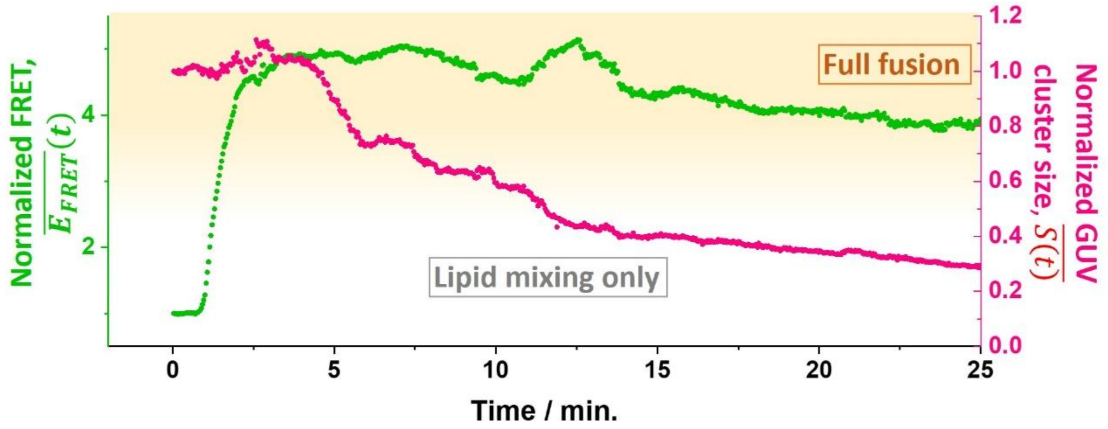


**Figure S7. Evolution of the normalized FRET signal and GUV cluster size following the addition of 5 mM $CaCl_2$ to 20:20:60 DOPS:DOPC:DOPE GUVs.** The normalized FRET signal $\overline{E_{FRET}}$ surpasses values measured in 20:80 POPS:POPS GUVs; compare to Fig. 3. The graph shows a rapid increase in lipid mixing due to the fusion cascade illustrated in Fig. 4 in the main text and Movie S2. The yellow shading marks conditions of fusion events became increasingly frequent. The fusion cascade results however in most of the GUVs bursting, leaving the microfluidic trap empty 15 minutes after the addition of calcium. While residual lipid in the trap continues to emit FRET signal and is still detected in the graph, the normalized cluster size, $\overline{S(t)} = \frac{S(t)}{S(t=0)}$ , decreases rapidly, capturing the loss of intact GUVs due to bursting.

### S8. FUSION PROCESS ARRESTING WITH HEMIFUSION

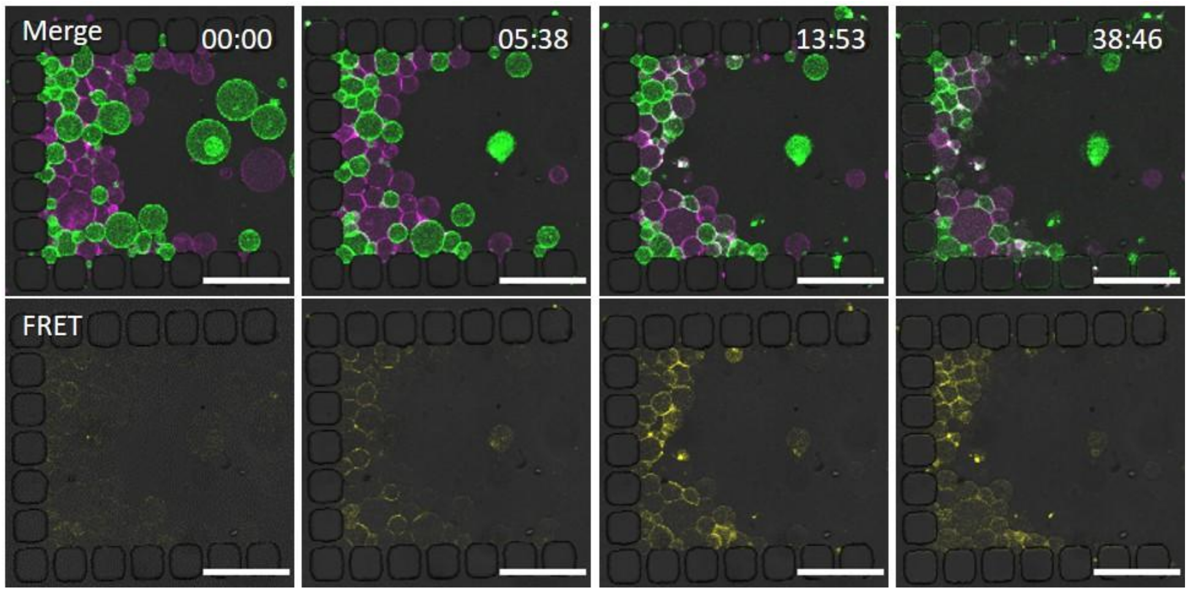


**Figure S8. 20:50:30 POPS:POPC:DOPE GUVs hemifuse when exposed to 10 mM [Ca2+], but they do not fuse.** The top images are overlay of bright-field and the two fluorescence channels (green and magenta) and the bottom images show an overlay of bright-field and FRET signal. Although after the addition of a calcium solution the GUVs adhere and start lipid mixing (as seen in the FRET channel), they do not fuse, even after 30 minutes of constant exposure to calcium. Time stamp format: mm:ss. Scale bars: 100 µm.

### S9. EXAMPLES OF FUSION AFTER THE ADDITION OF CALCIUM

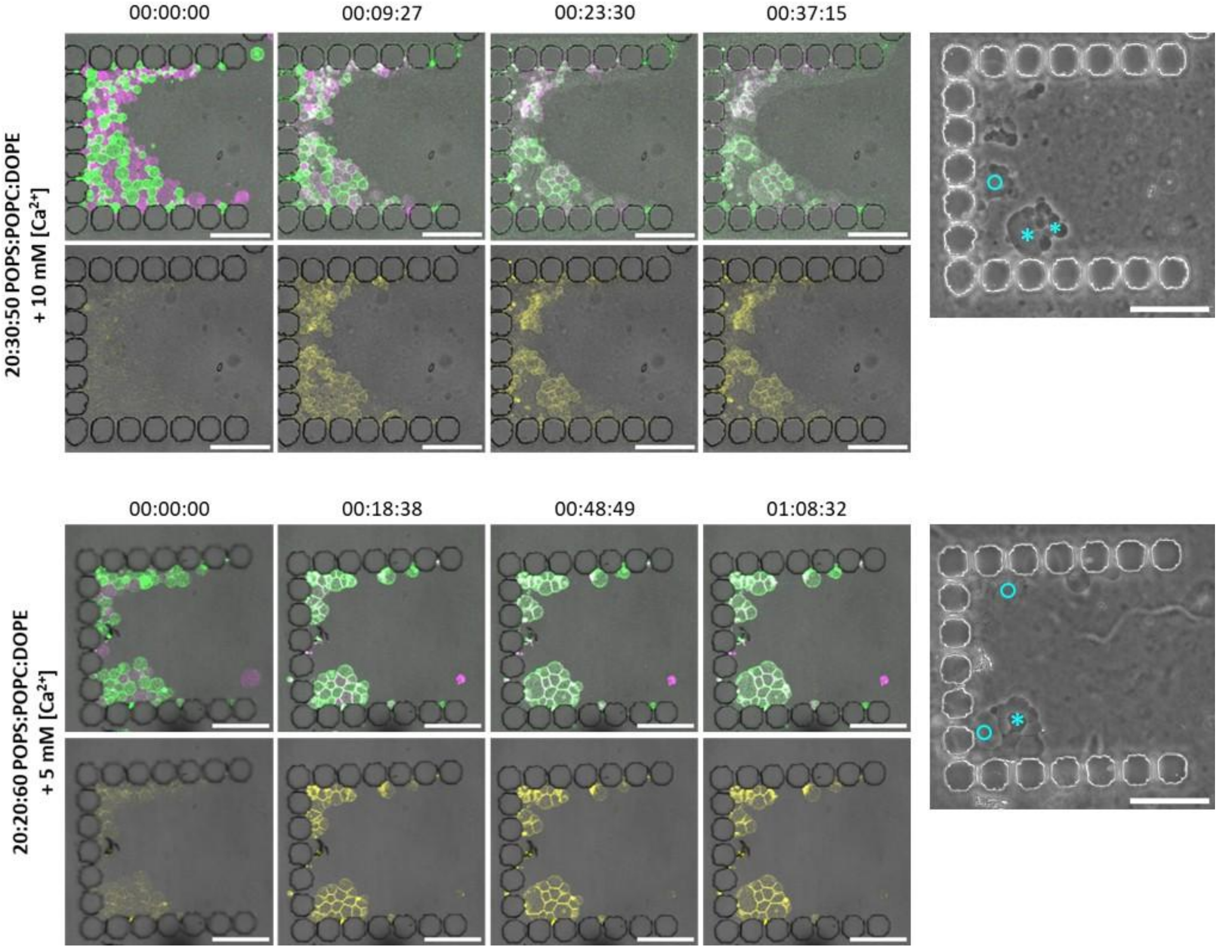


**Figure S9. Further fusion examples of GUVs with sufficiently high fraction of negative curvature lipids.** Top rows show Atto488 (green) and Atto633 (magenta) merged fluorescence and bright field signal, bottom – FRET channel overlaid with bright field signal. On the right side of the figure, the phase contrast images of the microfluidic traps at the end of the experiment show vesicles that have leaked (cyan circles) and intact vesicles (cyan asterisks). The 20:30:50 GUVs show pervasive bursting, however many fused vesicles remain intact long after the start of the experiment. Time stamp format: hh:mm:ss. Scale bars: 100 µm.

### S10. FRET EFFICIENCY OF INDIVIDUAL FUSION EVENTS

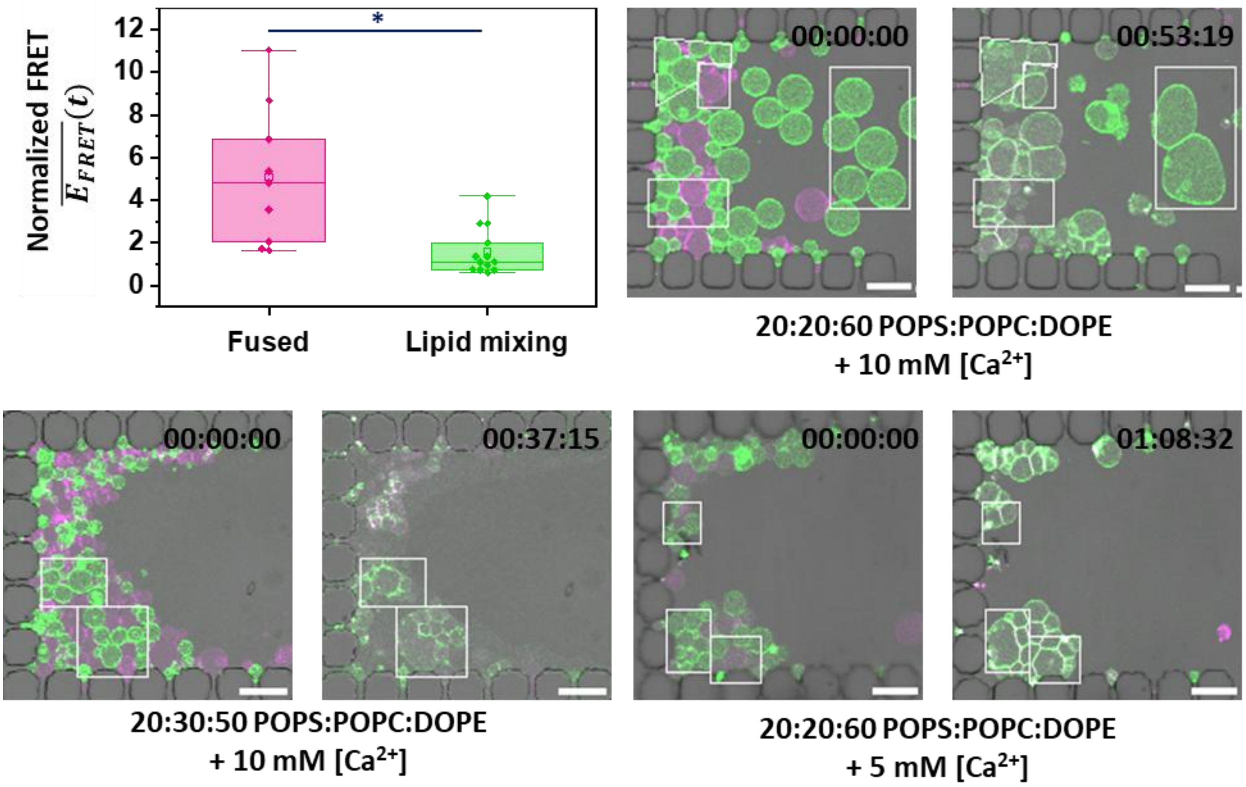


**Figure S10. Normalized FRET signal $\overline{E_{FRET}}(t)$ is consistently higher in fused GUVs compared to adhering or hemifused vesicles.** Confocal microscopy recordings were segmented (white frames) to track how FRET signal evolves in individual GUVs. Measurements were performed on 22 confocal microscopy segments. Box plots and statistical representation follow the conventions described in Fig. 2. Time stamp format: hh:mm:ss. Scale bars: 50 µm.

### S11. CLUSTER SIZE AND FRET SIGNAL AFTER FUSION

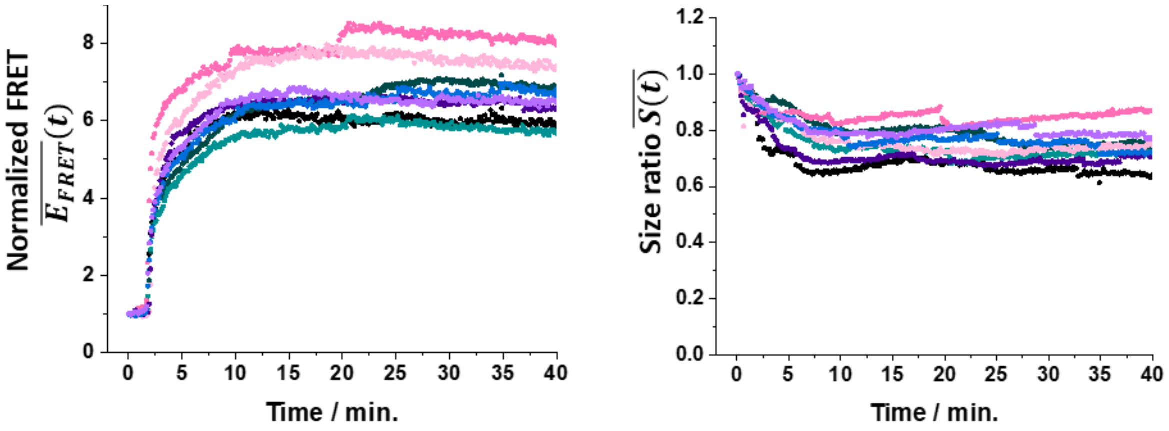


**Figure S11. Evolution of FRET and cluster size in fusogenic GUVs.** (A) Normalized FRET signal, $\overline{E_{FRET}}(t)$, for eight clusters of 20:20:60 POPS:POPC:DOPE GUVs following addition of 10 mM $Ca^{2+}$. (B) Corresponding normalized cluster size, $\overline{S(t)} = \frac{S(t)}{S(t=0)}$, where each color represents one cluster. Fusion occurs at different times and intensities depending on the initial cluster composition (see Figs. 3 and S1), making the kinetics highly situation-dependent. Despite sporadic fusion events and the apparent decrease in cluster size as vesicles attach and merge, the total cluster size decreases less dramatically than in cases of extensive bursting (Figs. 4 and S7), highlighting that full fusion can occur without widespread rupture.

### S12. EFFECT OF MEMBRANE CURATURE AND TENSION ON FUSOGENICITY

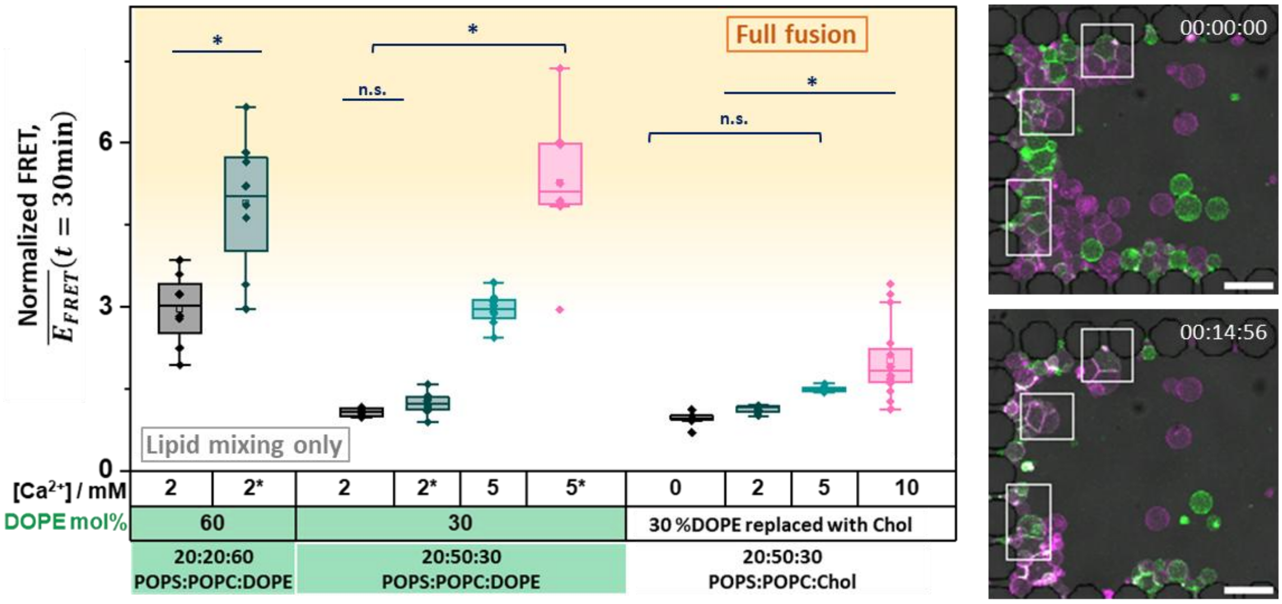


**Figure S12. Increased membrane tension grants higher fusion propensity only at high fractions of DOPE, while replacing DOPE with cholesterol does not increase fusion propensity.** The yellow-shaded region marks the conditions under which full fusion events became increasingly frequent, whereas conditions below this threshold led only to lipid mixing. The calcium concentrations marked with an asterisk are hypoosmolar solutions, which inflate the GUVs. Although membrane tension due to osmotic stress appears to increase lipid mixing, part of this effect can be attributed to pervasive GUVs bursting, with their membranes spreading and adhering to the neighboring GUVs. 88 GUV clusters were measured. Box plot representation and statistical analysis as described in Figs. 2 and 3. Right: example of hemifused 20:20:60 DOPS:DOPE GUVs before and after hypoosmotic treatment; despite some bursting, vesicles remain docked and interfaces straighten rather than relaxing to a spherical shape. Time stamp format: hh:mm:ss. Scale bars: 50 µm.

### S13. SUPPLEMENTARY MOVIES

All videos are available at https://owncloud.gwdg.de/index.php/s/I7obBx3whNMs4nz.

**Movie S1**. Confocal microscopy sequence of overlaid Atto488 (green), Atto633 (magenta) and bright-field channels showing the evolution of 20:80 POPS:POPC GUVs exposed to 10 mM $CaCl_2$ corresponding to the data shown in Fig. 3. No GUV-GUV fusion events occurred, only occasional vesicle rearrangement and disappearance of a GUV. Scale bar: 100 µm. Time stamp format: hh:ss:mm.

**Movie S2.** Confocal microscopy sequence of overlaid Atto488 (green), Atto633 (magenta) and bright-field channels showing the evolution of 20:20:60 DOPS:DOPC:DOPE GUVs exposed to 5 mM Ca $Cl_2$ corresponding to the data shown in Fig. 4 The GUVs rapidly undergo a cascade of fusion events, leading to the formation of large interconnected vesicles. This is followed by vesicle bursting, ultimately leaving the microfluidic trap nearly empty. Scale bar: 100 µm. Time stamp format: hh:ss:mm.

**Movie S3**. Confocal microscopy sequence overlaid Atto488 (green), Atto633 (magenta) and bright-field channels showing the evolution of 20:20:60 POPS:POPC:DOPE GUVs exposed to 10 mM $CalCl_2$ corresponding to the data shown in Fig. 6. The vesicles remain largely intact after fusion. Even 30 min after the onset of fusion, numerous fused GUVs persist within the trap, with minimal bursting. Scale bar: 100 µm. Time stamp format: hh:ss:mm.

**Movie S4**. Confocal microscopy sequence overlaid Atto488 (green), Atto633 (magenta) and bright-field channels showing the evolution to 20:50:30 POPS:POPC:DOPE GUVs exposed to 5 mM $CalCl_2$ and a hypotonic (257 mOsm/kg) solution. Some bursting is observed indicating that the osmolar gradient induces lysis, but no fusion events are observed. Scale bar: 100 µm. Time stamp format: hh:ss:mm.
